# The Necroptosis Effector MLKL drives Small Extracellular Vesicle Release and Tumour Growth in Glioblastoma

**DOI:** 10.1101/2021.01.12.426398

**Authors:** Gwennan André-Grégoire, Tiphaine Douanne, An Thys, Clément Maghe, Kathryn Jacobs, Cyndie Ballu, Kilian Trillet, Ignacio Busnelli, Vincent Hyenne, Jacky G Goetz, Nicolas Bidère, Julie Gavard

**Affiliations:** Team SOAP, CRCINA, Inserm UMR 1232, CNRS, Université de Nantes, Université d’Angers, Nantes, France; Institut de Cancérologie de l’Ouest (ICO), Saint-Herblain, France; INSERM UMR_S1109, Tumor Biomechanics, Université de Strasbourg, Fédération de Médecine Translationnelle de Strasbourg (FMTS), Strasbourg, France; CNRS SNC5055, Strasbourg, France

**Keywords:** Glioblastoma, MLKL, Extracellular Vesicles, Intracellular Trafficking, Temozolomide

## Abstract

Extracellular vesicles (EVs) are lipid-based nano-sized particles that convey biological material from donor to recipient cells. They play key roles in tumour progression, notably in glioblastoma in which the subpopulation of Glioblastoma Stem-like Cells (GSCs) might represent a meaningful source of tumour-derived EVs. However, the mechanisms involved in the production and release of EVs by GSCs are still poorly understood. Here, we report the identification of MLKL, a crucial effector of cell death by necroptosis, as a regulator of the constitutive secretion of small EVs from GSCs. The targeting of MLKL by genetic, protein depletion or chemical approaches alters endosomal trafficking and EV release and reduces GSC expansion *in vitro*. This function ascribed to MLKL appears independent of its role during necroptosis. *In vivo*, pharmacological inhibition of MLKL triggers a reduction of both the tumour burden in xenografted mice and of the level of plasmatic EVs. This work reinforces the idea of a non-deadly role for MLKL in endosomal trafficking and suggests that interfering with EV biogenesis is a promising therapeutic option to sensitize glioblastoma cells to death.

## Introduction

Glioblastoma (GBM) is the most prevalent and aggressive primary brain tumour in adults. This cancer remains mostly incurable with a median survival estimated at 14 months^1^. The therapeutic regimen, established in 2005^1^, relies on resective surgery, followed by concomitant radio-chemotherapy using the alkylating agent temozolomide (TMZ). Relapse is rapid and fatal, with scarce, palliative second line treatment options. Thus, optimized therapies are urgently needed. GBM aggressiveness partially relies on a subpopulation of rare, malignant cells with stem properties named GSCs (for *Glioblastoma Stem-like Cells*) that may act as a reservoir able to repeatedly initiate and repopulate the tumour mass^2^. GSCs also resist to radio and chemotherapy treatments^3,4,5^ and pervert the tumour microenvironment to support their maintenance, growth, and expansion.

Extracellular vesicles (EVs) are as important mediators of cell-to-cell communication within the tumour soil^6,7^. EVs are lipid-bilayer carriers, typically emanating from the intracellular endosomal compartment (exosomes) or budding at the plasma membrane (microvesicles). These nano-sized particles, ranging from 30–100 nm to few micrometres, haul proteins, lipids, and nucleic acids^8^. EVs are suspected to support cancer growth and dissemination through mediation of both local and at distance signalling^6,9^. For example, GBM-derived EVs have been shown to circulate in biofluids, and to transfer oncogenic material, such as the EGFRvIII variant^10^ or the active oncogene Ras^11^, to neighbouring non-tumour cells. Tumour EVs were also reported to assist in endothelium defects and angiogenesis. Indeed, the uptake by endothelial cells of tumour EVs, enriched with pro-angiogenic factors, results in the formation of tubules^12^ and the loss of the vascular barrier integrity^13,14^. Likewise, EVs contribute to cell proliferation, as well as tumour growth^12^, and are able to induce immunotolerance^15,16^. Alongside this role on the tumour stroma, EVs participate in GBM aggressiveness and resistance to treatments. For instance, TMZ impacts the number and content of EV-transported ribonucleic acids and proteins, which in turn convey TMZ resistance^5,17–20^. In this context, GSCs cope with TMZ toxic signals, while their protein cargoes shifted towards an adhesive signature^5^.

While virtually every cell type might empower the intracellular machinery to release EVs upon diverse stimuli^8^, cancer cells possess the privileged feature of constitutively hijacking this cell-to-cell communication route to dispatch mutations, metabolic changes, and resistance to treatments^12^. However, the detailed molecular mechanisms involved in tumour EV biogenesis and their further impact on disease progression remain unclear. In the search for novel mediators of constitutive EV release in cancer cells, clinical data mining identified here the Mixed Lineage Kinase domain-Like protein, MLKL, as of interest in GBM. This pseudokinase was initially identified as a key effector of necroptosis, a regulated form of necrosis occurring in apoptosis-deficient conditions^21,22^. However, MLKL also exerts non-necroptotic functions such as controlling adhesion molecule expression in endothelial cells^23^, regeneration after nerve injury^24^, NLRP3 inflammasome formation^25,26^, endosomal compartment trafficking and necroptosis-induced vesicles formation and extrusion^27–30,31^. MLKL therefore operates dual functions at the crossroads between live-death checkpoints and intercellular communication within the tumour microenvironment. By deploying a triad of complementary interfering approaches (siRNA, drug, and targeted protein degradation), we show that MLKL governs EV release and favours patient-derived GSCs *in vitro* and *in vivo*.

## Results

### Aggressive glioblastoma cells express MLKL

To identify Extracellular Vesicle (EV)-related genes whose expression might impact on the disease outcome, we interrogated The Cancer Genome Atlas (TCGA) online database through the Gliovis platform^32^. The analysis of 69 genes related to EV production^8^ and their hierarchization between high *versus* low expression, identified 9 candidates, which were significantly associated with probability of survival in GBM patients (Fig. 1a). Among these, the expression of *SYTL3, RAB27A* and *MLKL* were negatively correlated with glioblastoma patient survival (Fig. 1b). Besides its key function in orchestrating cell death via necroptosis^21,22^, MLKL was recently reported to play an important role in endosomal trafficking and EV biogenesis^27,28^. Alongside its prognosis value in survival probability, *MLKL* mRNA expression was heightened in GBM, as compared to non-tumour brains (Fig. 1c-d). This elevated expression was further confirmed at the protein level in brain biopsies (Fig. 1e). Of note, the level of *MLKL* mRNA is augmented according to the tumour grade (Fig. 1c). Likewise, MLKL expression is lower in patients with an isocitrate dehydrogenase *(IDH)-1* gene somatic mutation, typically associated with a younger age at diagnosis, and a better prognosis^33^ (Fig. 1c). Interestingly, *MLKL* expression was observed across the four reported molecular subtypes, with higher level in the most aggressive mesenchymal (MES) tumours^34^ (Fig. 1c).

**Figure 1.**
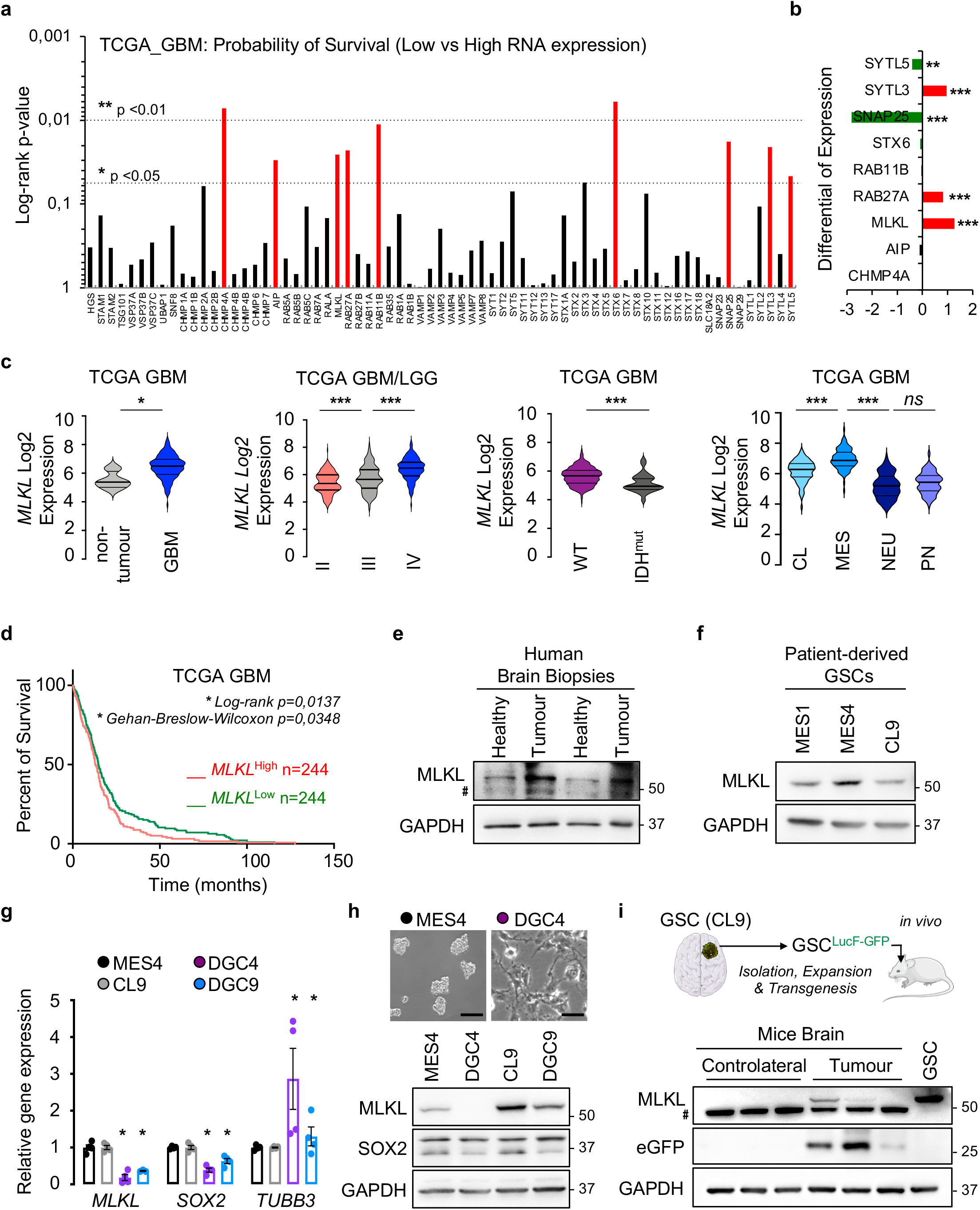
The expression of the pseudokinase MLKL is associated with aggressive glioma. **a** The Cancer Genome Atlas (TCGA RNAseq dataset) was interrogated *via* the GlioVis platform to analyse the probability of survival (log-rank p-value) in n=156 GBM patients, stratified based on low vs high expression (optimal cut-off) of 69 EV-related genes. **b** Fold changes in mRNA expression for genes selected from panel (a) in non-tumour (n=4) vs GBM patients, n=156. **c** Box plots of *MLKL* mRNA expression in non-tumour, low-grade glioma (LGG grades II and III) and in GBM (grade IV), according to the mutation status of *IDH1* (IDH1^mut^) and the GBM subtypes (classical CL, mesenchymal MES, neural NEU, and proneural PN), based on the TCGA RNAseq and Agilent4502A datasets. GBM status: non-tumour n=4, GBM n=156; Grades: II n=226, III n=244, IV n=150; IDH: WT n=339, mut n=27; Subtypes: CL n=86, MES n=96, NEU n=111, PN n=238. **d** Kaplan-Meier curve of the probability of survival for n=244 GBM patients with low vs high expression of *MLKL*, using optimal cut-off settings in the TCGA RNAseq dataset. **e** Protein lysates from tumour and control (Healthy) human brain tissue biopsies were analysed by immunoblot for MLKL protein expression. Hashtag symbol indicates non-specific signal. GAPDH serves as a loading control. **f** MLKL protein expression was assessed in three patient-derived Glioblastoma Stem-like Cells (GSCs) from mesenchymal (MES1 and MES4) and classical (CL9) subtypes. GAPDH serves as a loading control. **g** Histograms showing RNA level of *MLKL, SOX*2 and *TUBB3* genes by RT-qPCR analysis in GSCs (MES4 and CL9) and Differentiated Glioblastoma Cells (DGC4 and DGC9). Technical duplicates were performed in two independent experiments. **h** Representative bright field images of MES4 and DGC4. Scale bars: 100 μm and 30 μm. MLKL and SOX2 protein level was analysed in GSCs (MES4 and CL9) and DGCs (DGC4 and DGC9). GAPDH serves as a loading control. **i** Human GSCs (CL9) expressing eGFP-Luc were intracranially implanted in nude mice and protein samples harvested from xenografts were analysed for MLKL and eGFP expression by immunoblot. GAPDH serves as a loading control. Hashtag symbol indicates a non-specific signal. All immunoblots are representative of at least n=3. t-test and ANOVA, *p<0.05, ***p<0.001, and ns non-significant.

The expression of MLKL was next evaluated in patient-derived Glioblastoma Stem-like Cells (GSCs), a population of cells exhibiting stem-like properties thought to be responsible for initiation, maintenance and recurrence of the tumour^35,36^. This revealed that MLKL protein is readily detected in GSCs from classical (CL) and mesenchymal (MES) subtypes (Fig. 1f). Interestingly, the abundance of MLKL was found concomitantly reduced at the RNA and protein levels in Differentiated Glioblastoma Stem-like Cells (DGCs, Fig. 1g-h)^5^. Morphological differentiation, reduction in the expression of the SOX2 stem marker and increase in the expression of the TUBB3 neural differentiation marker confirmed their differentiation status (Fig. 1g-h). GSCs can also recapitulate tumour initiation when implanted in the striatum of immunocompromised mice (Fig. 1i). Protein extracts from such patient-derived xenografts retain a weak, but consistent level of expression of MLKL (Fig. 1i). Altogether, our data support the notion that MLKL expression is associated with aggressiveness in glioma clinical samples, as well as, *ex vivo* in patient-derived materials and, might notably be related to the stem-like cellular population.

### Glioblastoma Stem-like Cells are resistant to necroptosis

Because MLKL is a crucial effector of necroptosis^21,22^, we next investigated whether this form of cell death can occur in GSCs. Necroptosis can classically be triggered by stimulating cells with TNFα (T), while chemically obliterating both apoptosis and NF-κB-dependent pro-survival pathways with the pan-caspase inhibitor QVD (Q) and the SMAC-mimetic Birinapant (S), respectively^31^. Accordingly, in Jurkat T lymphocytes, TQS treatment caused a loss in phosphatidylserine asymmetry, together with a rupture of the plasma membrane, evidenced by Annexin V binding and PI incorporation, further culminating in cell death (Supplementary Fig. 1a-b). Likewise, the necroptosis inhibitor Necrostatin-1 (Nec-1) largely rescued Jurkat viability. In sharp contrast, three patient-derived GSCs previously described^36^, namely MES1, MES4 and CL9 GSCs, remained largely unaffected upon TQS challenge (Supplementary Fig. 1a-b). Yet, TNFα efficiently triggered the activation of the NF-κB pathway in GSCs, as illustrated by the phosphorylation of P65 and IκBα (Supplementary Fig. 1c). To explore in-depth the underlying mechanisms of tolerance towards necroptosis induction, the expression of receptor□interacting protein kinase□1 RIPK1 and RIPK3^22^, the upstream activators of MLKL, was monitored. Although RIPK1 was readily observed in GSCs, neither RIPK3 mRNA nor protein expression was detected in the three patient-derived GSCs tested (Supplementary Fig. 1d-e). In conclusion, our data unveils that the failure of patient-derived GSCs to complete the necroptotic program probably results from the lack of RIPK3 expression.

### MLKL controls the biogenesis of extracellular vesicles in glioblastoma stem-like cells

In addition to its role during necroptosis, MLKL functions in membrane trafficking and necroptosis-induced vesicle formation^27–29,31^. As EVs orchestrate GBM pathogenesis and aggressiveness^6^, we next explored the role of the pseudokinase in EV biogenesis in viable, proliferating mesenchymal GSCs. To this end, EVs were isolated from serum-free, exogenous EV-free cultured GSC supernatants by differential ultracentrifugation^37^, as depicted in Fig. 2a. As per the MISEV’s recommendations^38^, western-blot analysis validated the separation procedure, with the presence of two vesicular proteins, namely the transmembrane tetraspanin CD63 and the intravesicular accessory protein of the ESCRT machinery Alix, in the 100k fraction, together with the absence of the Golgi marker GM130 (Fig. 2b). Meanwhile, typical features of small extracellular vesicles were detectable in electron microscopy imaging (Fig. 2c). To assess the role of MLKL in EV biogenesis, the pseudokinase was first silenced via RNA interference (Supplementary Fig. 2a). We found that small EVs isolated in the 100k fractions of MLKL-silenced GSCs appear enlarged under electron microscopy (Fig. 2d). Single particle tracking analysis was deployed using Tunable Resistive Pulse Sensing (TRPS) technology (qNANO) to more accurately estimate both size and concentration of the EV preparations. This showed that MLKL silencing led to an overall reduction in the concentration of small EVs (100k pellets) released by GSCs, along with a modest increase in their mean size (Fig. 2e-f). As expected by the reported functions of RAB27A/B in vesicular trafficking^35^, a similar trend in EV concentration was observed in RAB27-silenced samples (Supplementary Fig. 2a-c). Of note, both quality and quantity of larger EVs (10k pellets) were not altered (Fig. 2e). This suggests that GSCs produced EVs with an increased diameter, yet less abundantly, when MLKL expression is reduced.

**Figure 2.**
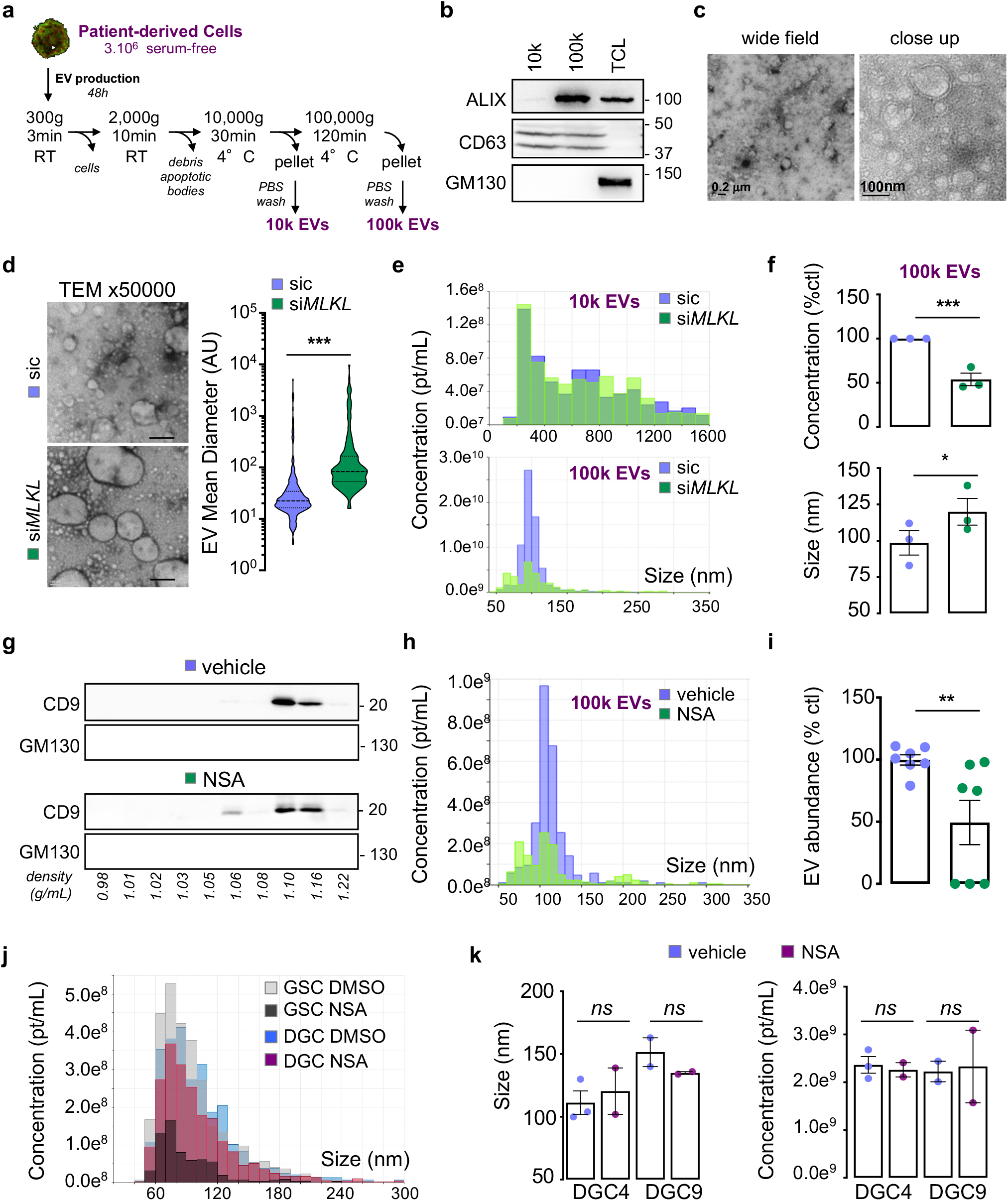
MLKL regulates extracellular vesicles released from glioblastoma stem-like cells. **a** Diagram of the isolation protocol for large (10k) and small (100k) Extracellular Vesicles (EVs) obtained from 48h-old conditioned media of serum-free, EV-free cultured Glioblastoma Stem-like Cells (GSCs). **b** Immunoblot analysis of 10k and 100k EV fractions isolated from mesenchymal GSCs (MES4) conditioned-media according to the protocol described in panel (a). Antibodies against EV specific markers (ALIX and CD63) and putative contaminant (GM130) were used as indicated. Total cell lysates (TCL) serve as an internal control. **c** Representative electron microscopy images of 100k EVs purified from MES4 conditioned-media according to the protocol described in panel (a), n=3. Scale bars 0.2 μm and 100 nm. **d** Representative electron microscopy images of 100k EVs from MES4 transfected with RNA duplexes targeting *MLKL* (si*MLKL*) and non-silencing duplexes (sic), imaged 72h. Scale bars: 300 nm. Quantification of the EV area observed (382 vs 153 EVs counted in two representative fields of view), n=3. **e** Representative diagrams of size distribution and particles concentration of large (10k) and small (100k) EVs isolated from sic and si*MLKL* transfected MES4, as estimated by tunable resistive pulse sensing analysis (TRPS, qNano, IZON). pt=particle, n=3. **f** Histograms quantifying relative particle concentration (% of control) and mean size diameter of small EVs (100k) shown in panel (e), n=3. **g** Density gradient analysis by immunoblot of 100k EVs for a specific EV marker (CD9) and putative contaminant (GM130) in the 10 collected fractions, labelled from top to bottom. EVs were isolated from MES4 treated for 48h with vehicle (DMSO) and the MLKL inhibitor NSA (5 μM), n=2. **h** Representative diagrams of size distribution and particle concentration of small EVs (100k) in vehicle (DMSO) and NSA-treated MES4, as measured with TRPS. pt=particle, n=2. **i** Quantification by ELISA of 100k EVs isolated from 48h-old vehicle and NSA-treated MES4, n=7. **j** Representative diagrams of size distribution and particle concentration of small (100k) EVs in vehicle and NSA-treated MES4 and Differentiated Glioblastoma Cells (DGCs, DGC4), obtained by TRPS. pt=particle. **k** Histograms quantifying particle concentration and mean size diameter of small EVs (100k) shown in panel (j). pt=particle, n>2. All immunoblots are representative of at least n=3, unless specified. Mann-Whitney test, **p<0.01, ***p<0.001 and ns not-significant.

To confirm our findings pharmacologically, we blocked MLKL with the known inhibitor necrosulfonamide (NSA)^22^. Separation by density gradient, followed by immunoblotting analysis, for tetraspanin CD9 revealed that EV fractions floated at an expected density of around 1.10 g/mL (Fig. 2g). However, a fraction of CD9-positive particles emerged at lower density when cells received the MLKL inhibitor NSA (Fig. 2g), suggesting a shift in vesicle size upon treatment. Likewise, TRPS measurements unmasked the generation of higher diameter vesicles in NSA-challenged GSCs (Fig. 2h). This was again accompanied by a decrease in EV abundance (Fig. 2h-i), thus mirroring *MLKL* siRNA-provoked phenotype. Lastly, EVs emanating from differentiated glioblastoma cells (DGCs), in which MLKL expression is strongly reduced (Fig. 1g-h) remained largely unaffected by MLKL pharmacological inhibition with NSA, in terms of size and concentration (Fig. 2j-k). Overall, our data reinforce the idea of a routine function of MLKL in the biogenesis of EVs in GSCs.

To gain further insights in how MLKL controls EV release, we next carried out a mass-spectrometry-based comparative analysis of the protein cargoes in EVs isolated from naive and NSA-challenged GSCs (Fig. 3a, Supplementary Table 1, 48h treatment). Interestingly, when analysing known and predicted protein-protein interaction networks, as well as enriched pathways using the STRING open data source, NSA treatment steered the content of EVs towards proteins involved in biological processes related to transport and localisation, suggesting that MLKL inhibition might relate to EV processing and/or release (Fig. 3a). This putative function was further substantiated with a transcriptomic analysis of cellular RNA upon NSA challenge (at 16h, and 48h, Fig. 3b, left panel). Principal component analysis shows that the first variance can be explained by the duration of NSA exposure (early time point on the left 16h *vs* 48h), while second variance discriminated between untreated and treated samples (DMSO above *vs* NSA bottom). Differentially expressed genes (q-value) were then hierarchized based on their Gene Ontology, unmasking again an organelle trafficking signature, with some of the most typical genes highlighted on the volcano plot (Fig. 3b, middle and right panels) (Supplementary Table 2).

**Figure 3.**
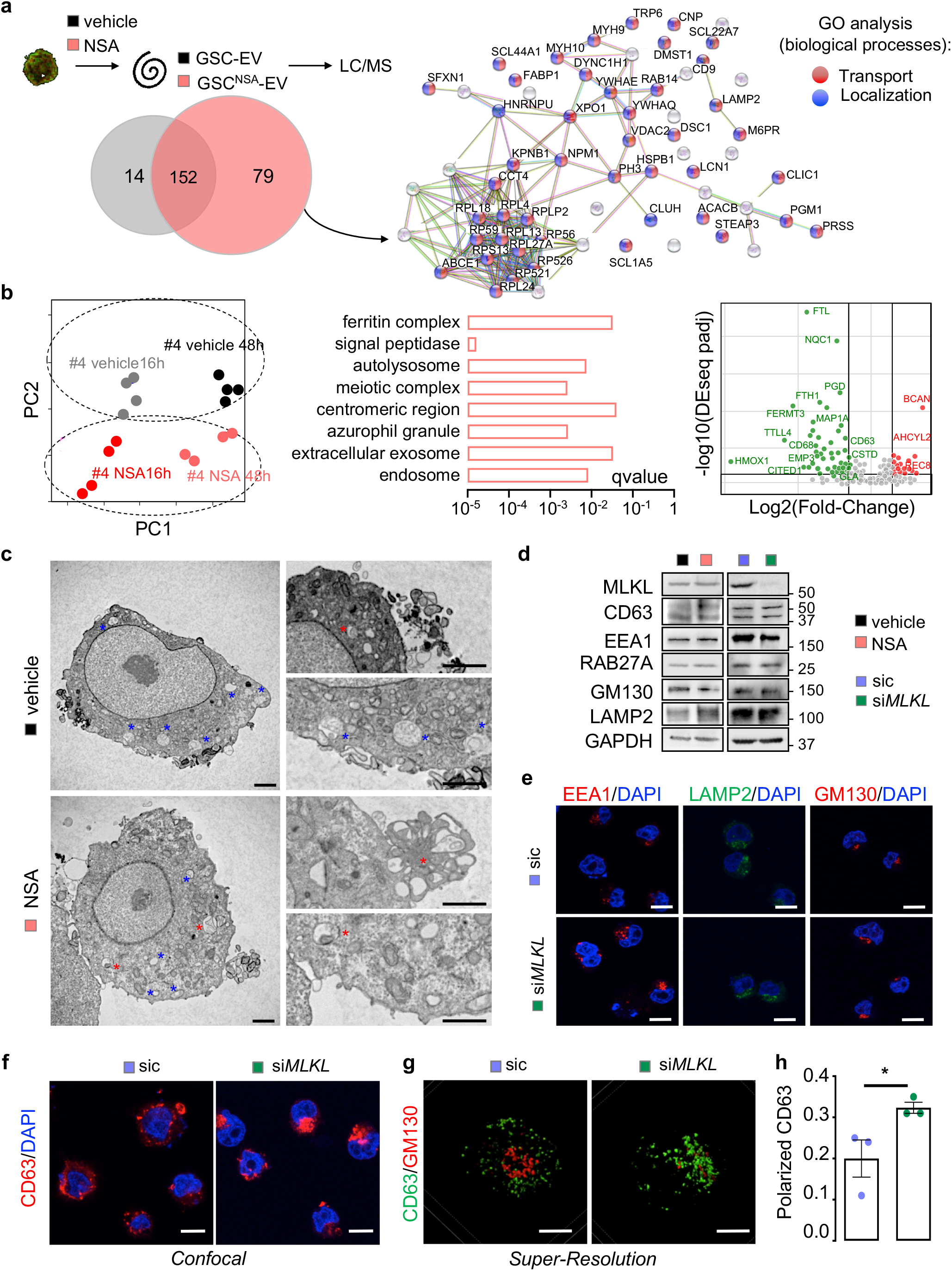
MLKL inhibition impairs constitutive biogenesis of extracellular vesicles in glioblastoma stem-like cells. **a** Mass spectrometry analysis (LC-MS/MS) of the protein content of small Extracellular Vesicles (EVs) isolated from mesenchymal Glioblastoma Stem-like Cells (MES4 GSCs) treated with vehicle (DMSO) and the MLKL inhibitor NSA (5 μM) for 48h. Right panel shows STRING analysis of the 79 proteins detected only in EVs isolated from NSA-treated cells. Main gene ontology (GO) biological processes are highlighted in red (Transport) and blue (Localization). **b** Transcriptomic analysis by 3’Sequencing RNA Profiling of MES4 treated with vehicle (DMSO) and NSA (5 μM) for 16h and 48h. Left panel represents the principal component analysis of quadruplicates. Middle panel, analysis of GO terms over- or under-represented. Right panel, specific genes under- (green) or over- (red) expressed in NSA-treated MES4 cells. **c** Representative electron microscopy images of vehicle (DMSO) and NSA-treated MES4 for 48h. Blue stars pointed to multivesicular bodies (MVB), red stars abnormal enlarged MVB. Scale bars: 2 and 1 μm. **d** MES4 were either challenged with DMSO and NSA for 48h, or transfected with non-silencing (sic) and MLKL targeting RNA duplexes (si*MLKL*) for 72h prior analyses of organelles and EV-biogenesis proteins by immunoblot. GAPDH serves as a loading control. **e-f** Representative confocal images of MES4 transfected with sic and si*MLKL* for 72h, fixed and immunostained as indicated. Nuclei were counterstained with DAPI (blue). Scale bars: 10 μm, n=3. **g** Representative super-resolution structured illumination microscopy (SIM) was performed to analyse CD63 distribution in sic and si*MLKL* transfected MES4. GM130 stained Golgi. Scale bars: 4 μm, n=2. **h** Polarized immunostaining of CD63 was counted from panel (f) (n>10 per sample, in three independent experiments). All immunoblots are representative of at least n=3, unless specified. Mann-Whitney test, *p<0.05.

In addition to RNA and protein changes, electron microscopy images uncovered large structures budding at the plasma membrane upon NSA treatment, reminiscent of ghostlike multi-vesicular bodies (MVB)^28^ (Fig. 3c). No major action of NSA was detected on both the level of expression and the subcellular localisation of intracellular organelle components, including endosomes, lysosomes, and Golgi (Fig. 3d). Similar results were obtained when *MLKL* was silenced (Fig. 3d-e). However, both confocal and super-resolution microscopy unveiled re-localization and polarization of CD63 staining, a classical tetraspanin marker for EVs enriched in late endosomes and lysosomes^39^, in GSCs knocked down for *MLKL* (Fig. 3f-h). Our data thus point towards a defect in MVB subcellular trafficking, orchestrated by MLKL in these glioblastoma cells.

### Interfering with MLKL reduces GSC expansion

To evaluate the functional impact of interfering with MLKL-dependent EV release in GSCs, we first deployed a siRNA-based strategy. We observed a reduction in the size and number of tumour spheroids from *MLKL*-silenced cells, although the level of expression of the classical markers for stemness NESTIN and SOX2 were not overtly changed (Fig. 4a-c). Because the impairment of tumoursphere formation might be linked to proliferation, EdU incorporation was tracked in *MLKL* siRNA-transfected GSCs. Both confocal images and flow cytometry demonstrated that proliferation was reduced in *MLKL* knocked down cells (Fig. 4d-f). This was accompanied by a flip of phosphatidylserine, indicating that viability was blunted (Fig. 4g-h). In agreement with this, MLKL silencing coincided with a concomitant increase in apoptosis and a decrease in cell cycle progression through mitosis, as detected via Caspase 3 cleavage and histone H3 phosphorylation, respectively (Fig. 4i). Real-time, automated imaging confirms the impairment in cell viability upon NSA treatment (up to 4 days), resulting in smaller tumourspheres (Supplementary Movie 1). Of note, this phenotype did not appear to rely on a loss of EV uptake in GSCs, as 10k and 100k EVs were equally internalized in control and *MLKL*-silenced GSCs (Supplementary Fig. 3). Additionally, *MLKL* knockdown mimicked the silencing of *RAB27A/B* secretory proteins in terms of viability and cell death (Supplementary Fig. 2d-e). In order to eliminate directly and acutely the pool of translated proteins, MLKL protein was depleted using antibody-targeting degradation, by deploying the trim-away approach^40^ (Fig. 4j). Flow cytometry analysis of cell death mimicked the results obtained with RNA interference, as the proportion of PI-incorporated dead cells was potently augmented when MLKL protein was reduced (Fig. 4k). Finally, a pharmacological approach was employed using NSA. Recapitulating RNA and protein interference data, NSA administration hindered spheroid size and formation (Fig. 5a-b). Self-renewal ability was further halted as estimated with limiting dilution assay and stem cell frequency index (Fig. 5c). Viability was reduced in NSA conditions, in GSCs but not in their DGC sister cells that express lower amount of MLKL (Fig. 5d). Mirroring siRNA experiments again, proliferation was reduced and cell death increased upon MLKL inhibition (Fig. 5e-g). The alkylating agent temozolomide (TMZ) is the standard-of-care chemotherapeutic drug to which GSCs are largely resistant *in vitro* and *in vivo*^5^. While TMZ induced the phosphorylation of ATM^41^, blocking MLKL did not alter TMZ signalling to ATM (Fig. 5h). Interestingly, cell death was strengthened upon TMZ challenge in *MLKL*-silenced cells (Fig. 5i). Accordingly, viability declined when MLKL inhibition was combined with TMZ administration (Fig. 5j), suggesting a possible additive and/or permissive action of MLKL blockade on TMZ challenge. Thus, blocking MLKL opposed GSC expansion *in vitro*.

**Figure 4.**
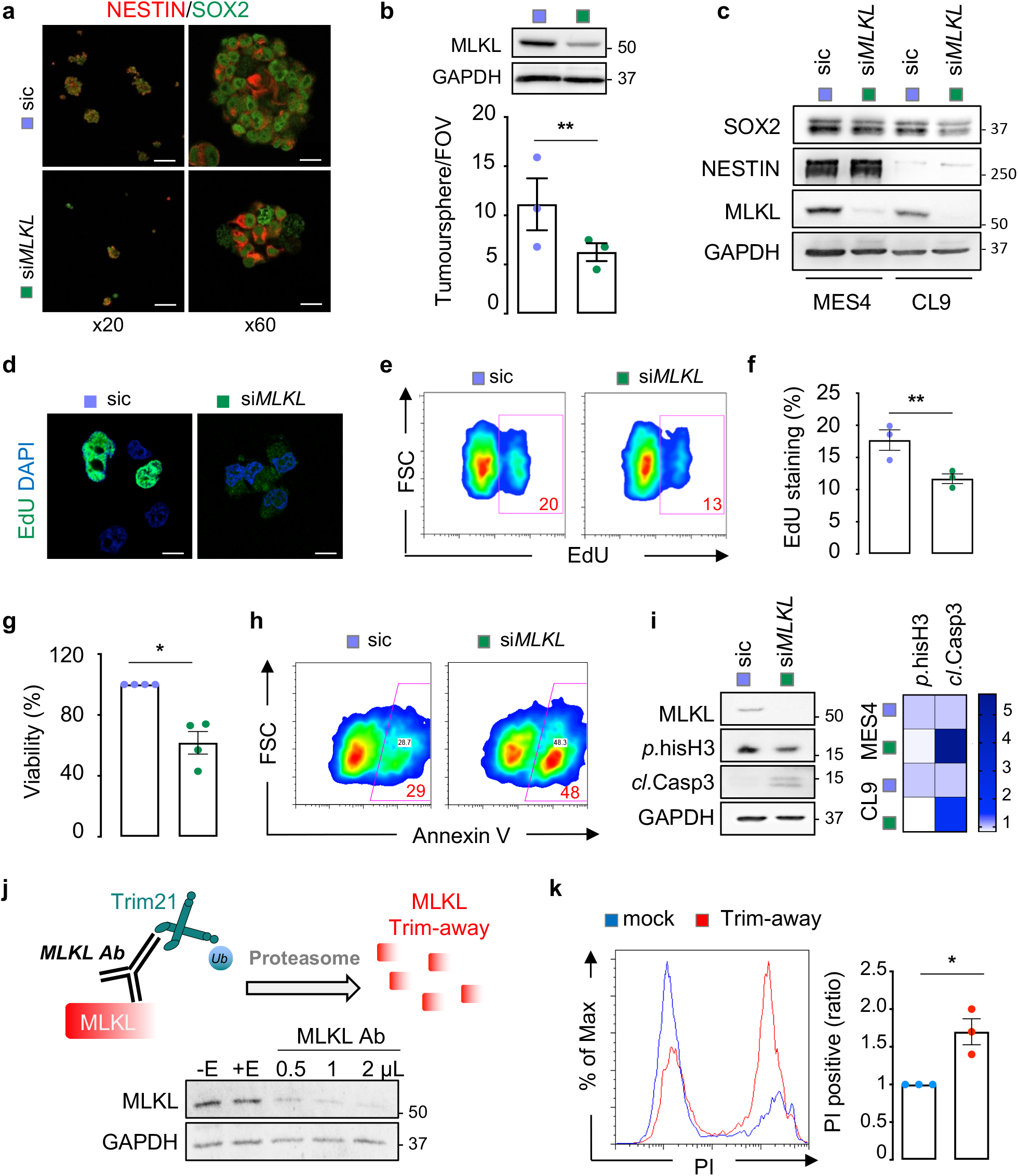
MLKL knockdown reduces glioblastoma stem-like cell expansion *in vitro*. **a** Mesenchymal Glioblastoma-Stem like Cells (MES4 GSCs) transfected with non-silencing (sic) and MLKL targeting RNA duplexes (si*MLKL*) were analysed after 72h. Representative confocal images of the stemness markers NESTIN (red) and SOX2 (green), as indicated. Scale bar x20: 100µm; x60: 16 µm. **b** Upper panel, knockdown efficiency of *MLKL* in sic and si*MLKL* transfected MES4 was verified after 72h by immunoblotting of MLKL. GAPDH serves as a loading control. Lower panel, tumoursphere formation assay (manually counted per field of view, FOV) for 4 days in sic and si*MLKL* transfected MES4. **c** The expression of SOX2 and NESTIN was evaluated by immunoblot in MES4 and CL9 cells transfected with sic and si*MLKL* for 72h. GAPDH serves as a loading control. **d** Representative confocal images of EdU incorporation (green) in sic and si*MLKL* MES4. Nuclei were counterstained with DAPI (blue). Scale bars: 10 µm. **e** Representative flow cytometry analysis of EdU incorporation in sic and si*MLKL* MES4. **f** The percentage of EdU-positive cells was quantified from panel (e). **g** Uptiblue viability assay in sic and si*MLKL* MES4 at 48h. **h** Representative flow cytometry analysis of the cell death marker Annexin V surface staining sic and si*MLKL* MES4 at 72h. **i** Protein lysates from sic and si*MLKL* MES4 and classical GSCs (CL9) were analysed by immunoblot for proliferation and death with phospho-Histone H3 (*p*.hisH3) and Cleaved Caspase-3 (*cl*.Casp3), respectively, and further quantified by densitometry (right panel), n=2. MLKL knockdown was verified. GAPDH serves as a loading control. **j** Diagram of endogenous MLKL protein was directly targeted for degradation by Trim-away technique using MLKL antibody (Ab). MLKL levels were analysed in MES4 by immunoblot 16h after electroporation of increasing Ab quantities. GAPDH serves as a loading control. “E” means electroporation. **k** Quantification by flow cytometry of propidium iodide (PI) incorporation in MES4 after MLKL trim-away overnight (16h). Histograms showing the ratio between PI-positive and total cells. Data are representative of at least three independent experiments, unlike otherwise stated. Mann-Whitney test, *p<0.05, **p<0.01.

**Figure 5.**
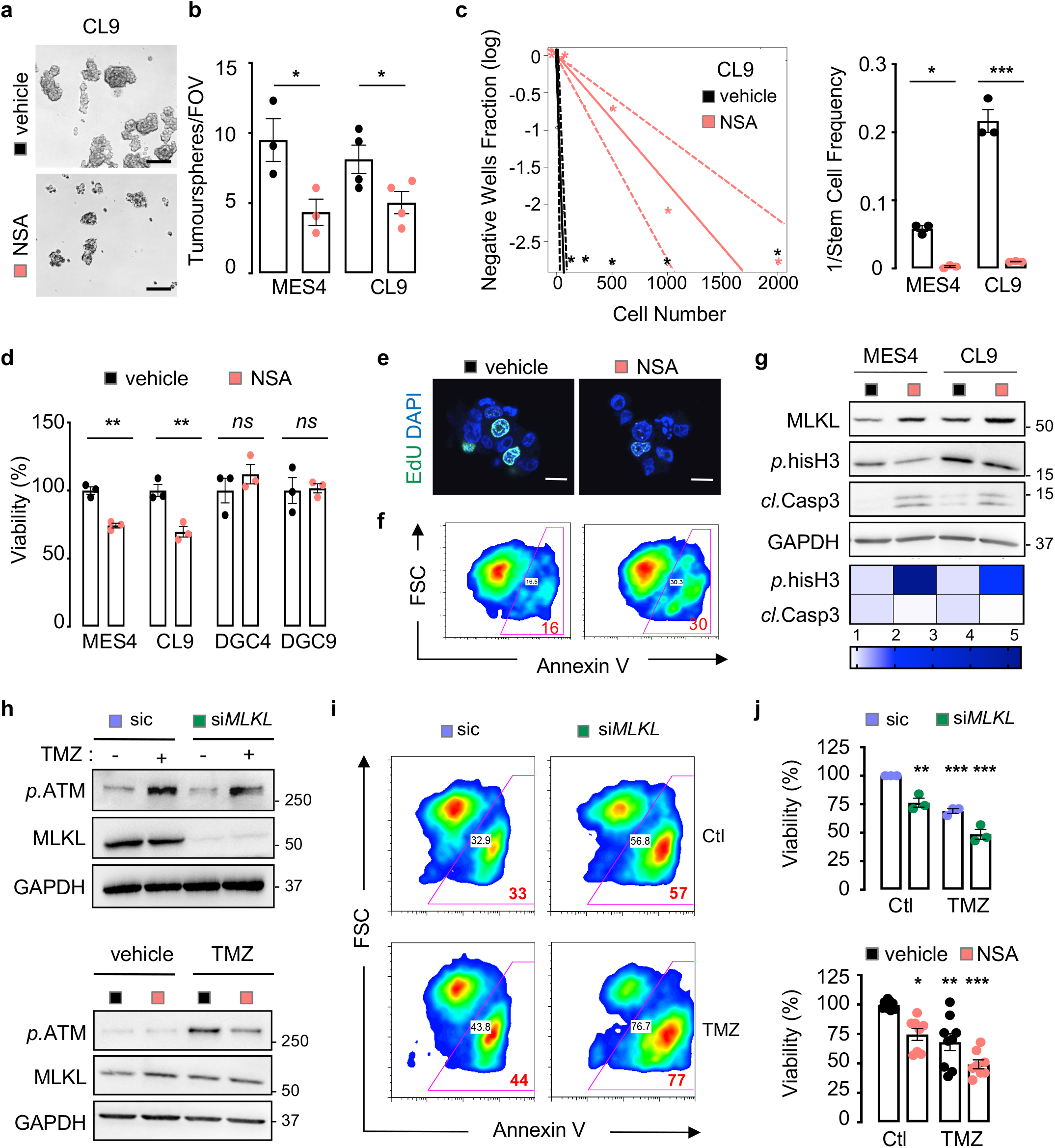
Pharmacological inhibition of MLKL impairs GSC expansion and potentialize the standard of care *in vitro*. **a** Representative bright field images of classical Glioblastoma Stem-like Cells (CL9 GSCs) treated with vehicle (DMSO) or the MLKL inhibitor NSA (5 μM) for 48h. Scale bars: 100 μm. **b** Tumoursphere formation assay (manually counted per field of view, FOV) was performed in mesenchymal GSCs (MES4) and CL9 cells in response to vehicle and NSA treatments for 4 days. **c** Limiting dilution assay (LDA) of CL9 in response to vehicle and NSA for 14 days. Left panel, representative linear regression plot for DMSO and NSA treated CL9 cells. Right panel, LDA estimation of the stem cell frequency in MES4 and CL9. **d** CellTiter-Glo viability assay of GSCs (MES4 and CL9) and corresponding Differentiated Glioblastoma Cells (DGC4 and DGC9) after 48h treatment with vehicle and NSA, n=3. **e** Representative confocal images of proliferation assessed with EdU incorporation (green) in MES4 after 48h treatments with vehicle and NSA. Nuclei were counterstained with DAPI (blue). Scale bars: 10 μm. **f** Representative flow cytometry analysis of the cell death marker Annexin V surface staining in MES4 after 48h treatment with vehicle and NSA, n=3. **g** Protein lysates from vehicle and NSA treated MES4 and CL9 were analysed by immunoblot for proliferation and death with phospho-Histone H3 (*p*.hisH3) and Cleaved Caspase-3 (*cl*.Casp3), respectively, and further quantified by densitometry (bottom panel), n=2. MLKL and GAPDH serve as loading controls. **h** Immunoblot analysis of the phosphorylation of ATM (alkylation signature, *p*.ATM) in MES4 treated with the standard-of-care Temozolomide (TMZ, 100 µM) with *MLKL* silencing and MLKL inhibition (NSA). MLKL and GAPDH serve as internal and loading controls. **i** Representative flow cytometry analysis of the cell death marker Annexin V surface staining in MES4 treated as in panel (h). **j** Uptiblue viability assay after 48h in MES4 treated as in panel (h), n=8. Data are representative of at least three independent experiments, unlike otherwise stated. Mann-Whitney test and one-way ANOVA, *p<0.05, **p<0.01, ***p<0.001.

### Pharmacological targeting of MLKL *in vivo* inhibits glioblastoma tumour growth

Our results prompted us to evaluate the effect of MLKL inhibition in an orthotopic *in vivo* model of glioblastoma in nude mice (Supplementary Fig. 4a-e)^36,42^. First, to determine the bio-safety of NSA, mice were administered 5 mg/kg of NSA tri-weekly for 4 weeks by intra-peritoneal route (Fig. 6a). Gross animal examination did not reveal alterations of the general health status between healthy animals that received either NSA or vehicle (Fig. 6b). The same was true with complete blood count (Supplementary Table 3). While the weights of the heart and kidneys did not seem to suffer from NSA challenge, the livers appear slimmer, although plasmatic levels of ASAT and ALAT were unchanged, overall suggesting no overt adverse effects of NSA administration *in vivo* (Fig. 6c-d).

**Figure 6.**
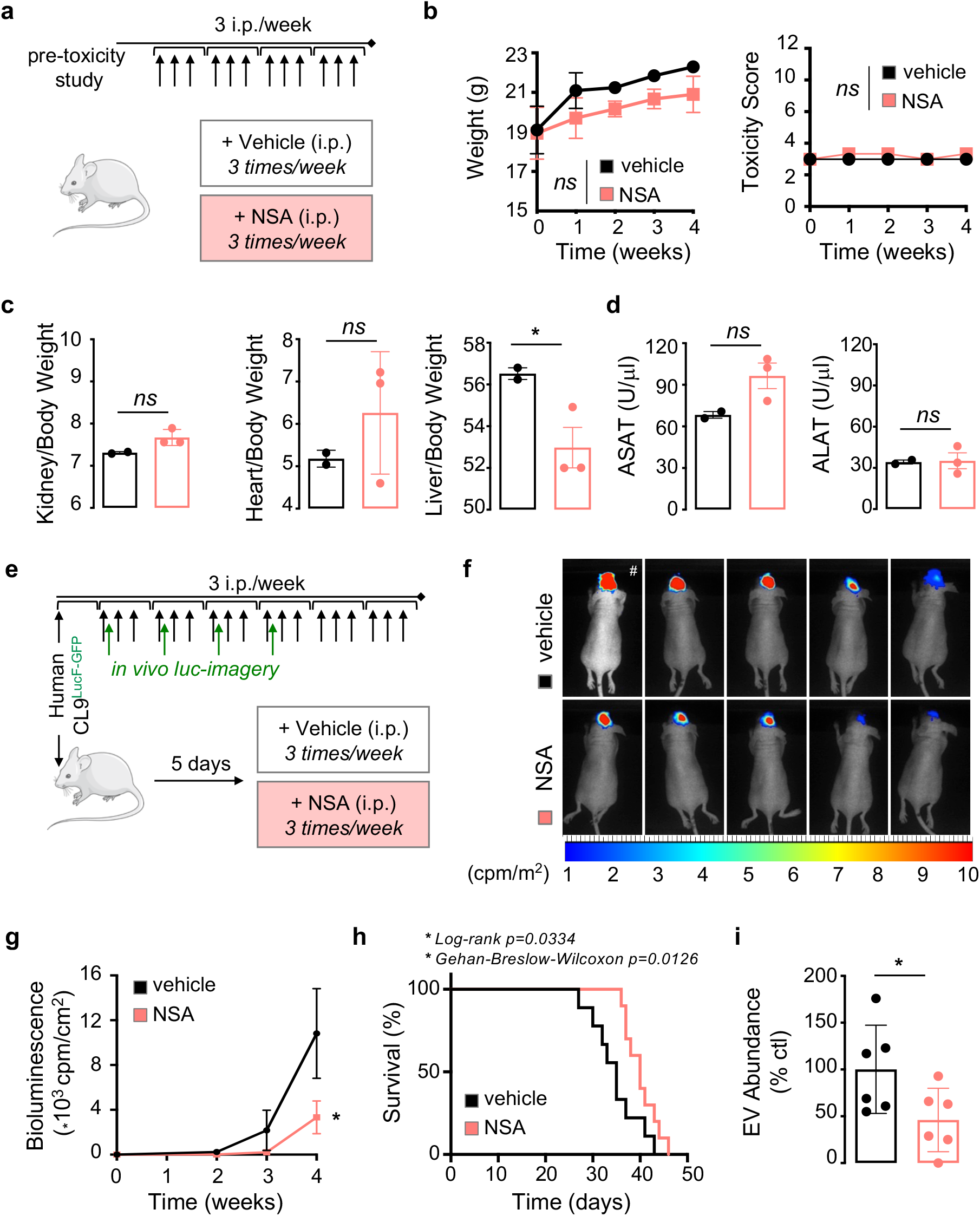
NSA administration slows down tumour growth *in vivo*. **a** Diagram of protocol for evaluation of NSA tolerance/toxicity *in vivo* in mice. **b** Follow-up of the body weight (left panel) and behaviour and general health parameters (toxicity score, right panel) in tumour-free mice treated with vehicle (DMSO) and NSA for 4 weeks (IP 5 mg/kg, 3 times/week), n=2 (vehicle) and 3 (NSA). **c** Kidney, heart and liver were observed and weighted at sacrifice (week 4) for toxicity assessment, n=2 (vehicle) and 3 (NSA). **d** Liver function assessment using the transaminase ASAT and ALAT measurements in plasma from vehicle and NSA treated mice for 4 weeks, n=2 (vehicle) and 3 (NSA). **e** Diagram of implantation of patient-derived classical Glioblastoma Stem-like Cells (CL9 GSCs, expressing luciferase and GFP), and treatment regime with vehicle and NSA IP 5 mg/kg, 3 times/week). Follow-up of tumour growth is performed via luciferase (luc)-based live imaging. **f** Representative bioluminescence images of brain tumours at week 4 (except #, done at week 3 before death), n=5. **g** Quantification of luminescence (count per minute, cpm/cm^2^) in mice treated with vehicle and NSA, as depicted in panel (e), n=5. **h** Kaplan-Meier survival curve of mice treated with vehicle and NSA, n=10 from two independent experiments with five mice. **i** Quantification by CD63-ELISA of vesiclemia, EV abundance of circulating 100k small EVs, isolated from plasma of vehicle (DMSO, black) and NSA (pink) treated mice. Data are expressed as the percentage of vehicle, n=6, from two independent experiments. Data are representative of at least three independent experiments, unlike otherwise stated. Mann-Whitney test and two-way ANOVA, *p<0.05 and ns not-significant.

In order to gain further insights into the therapeutic potential of MLKL inhibition in glioblastoma, nude mice were implanted with GFP-Luc-expressing CL9 GSCs into the striatum and treated with NSA (5 mg/kg) three times a week, starting one week post-grafting in randomized animals (Fig. 6e). Tumour progression was monitored with bioluminescence imaging once a week together with signs of morbidity (Fig. 6e). Tumour-emanating bioluminescence was quantitatively reduced upon NSA intraperitoneal administration (Fig. 6f-g). Importantly, NSA injection significantly improved the overall survival of tumour-bearing mice compared to their vehicle-treated counterparts (Fig. 6h), suggesting an overall reduction in tumour burden. In order to control whether this beneficial action was linked to EV release, murine blood was collected at sacrifice and circulating EVs were isolated. In keeping with the idea that vesiclemia, *i*.*e*. the concentration of EVs in plasma, correlated with tumour evolution^43^, the abundance of circulating EVs was dampened in animals that received NSA (Fig. 6i). Collectively, these *in vivo* data provide a strong basis for an instrumental role of MLKL in the production of tumour-derived EVs.

## Discussion

Extracellular vesicles operate as instrumental conveyors in cancer settlement and progression^44^. They orchestrate the delivery of tumour-emanating signalling cues in the local environment and at distance, and therefore contribute to diseases. Likewise, glioblastoma cells constitutively release large amount of EVs, playing important roles in gliomagenesis, response to treatments and cell-to-cell communication within the tumour soil and response to treatments^5,10–20^. In this context, interfering with EV-based communication might impede tumour growth. However, the direct impact of blocking EV biogenesis in the source cell is not fully elucidated. Here, we provide the proof-of-concept that targeting MLKL-dependent release of glioblastoma cell-produced EVs is associated with a reduction in glioblastoma cell growth.

During necroptotic cell death, MLKL phosphorylation by RIPK3 drives its oligomerization and insertion in the plasma membrane, thereby forming lytic pores^22,31,45,46^. Although several compounds were shown to trigger necroptosis *in vitro* in glioblastoma cell lines^47,48,49,50^, our data show that this is not the case with patient-derived GSCs most likely due to the absence of RIPK3. This fits with the idea that RIPK3 expression is silent in most cancer cells and tissues due to epigenetic changes^51,52,53^. Indeed, the down-regulation of RIPK3 expression owing to genomic methylation of the RIPK3 promoter blocks, at least partially, necroptosis-induced cell death by several chemotherapeutic agents^51,52,53^. The low expression of RIPK3 is associated with a poor prognosis in oesophageal squamous cell carcinoma^54^, while RIPK3 status was recently proposed as prognostic marker for low grade glioma^55^. Thus, GSCs might hijack epigenetic regulation of RIPK3 gene expression to shield against MLKL-dependent necroptosis cell death.

In viable cells, MLKL was shown to undergo phosphorylation, independent of RIPK3, and to associate with intracellular vesicles, where it regulates constitutive endosomal trafficking and the extrusion of EVs^28^. Whether MLKL undergoes phosphorylation and oligomerization to regulate EV genesis in GSCs is still under investigation. Our results with NSA, which prevents MLKL multimerization via covalent interaction with human MLKL residue Cys86, while sparing its phosphorylation, suggest that similar mechanisms might be at play in GSCs. Multiomics analysis of EV content, as well as a detailed look at the the global structure of organelles, suggest that the loss of MLKL function corroborates with defective multivesicular body (MVB) maturation route. This step is described to require the RAB27A/B small GTPases, which facilitate the trafficking of multivesicular bodies (MVB) towards the cell periphery, as well as their docking and fusion to the plasma membrane^8,31,39^. MLKL interference phenocopies RAB27A/B silencing in patient-derived cells, in terms of EV release and cell viability^53,57^. Likewise, altering RAB27A/B functions was shown to reduce migration and invasiveness in a model of glioma cell line, by manipulating lysosomal cathepsin D exocytosis^58^. Although lysosomes do not seem to be overtly affected upon MLKL inhibition in GSCs, further studies will decipher whether MLKL blockade shapes lysosome functions, an identified vulnerability point in glioblastoma^59,60^.

In keeping with the idea of limiting EV-dependent communication within the tumour microenvironment, MLKL might emerge as a potential druggable target. First, *mlkl knockout mice* are viable without obvious abnormalities, suggesting therefore, that obliterating MLKL *in vivo* does not exert major toxicity^61^. The maximal nontoxic dose of NSA injected intraperitoneally in mice was determined at 20 mg/kg^62^. In addition, the small size of sulfonamide compounds suggest that NSA may cross the blood-brain-barrier in mice even when administered via systemic routes^63^. Here, we used a chimeric model in which human patient-derived cells are transplanted into immunosuppressed mouse brains. Because NSA targets human, but not murine MLKL protein^22^, NSA might impair directly human tumour cell growth within the brain, reinforcing the idea that NSA crosses the blood-brain-barrier in the xenograft models. Of note, NSA challenge not only reduced the concentration of circulating EVs in the peripheral blood of intracranial tumour-bearing mice, but also tumour burden, culminating in the extension of survival. Targeting EVs *in vivo* to reduce tumour growth has already been proposed. For example in GBM, miR-1 overexpression in GSCs was able to modify EV protein cargoes and to ultimately reduce tumourigenicity, invasiveness and angiogenesis *in vivo* in xenografted mice^64^. In addition *RAB27A* silencing resulted in decreased primary tumour growth and lung dissemination of metastatic carcinoma^56^. Furthermore, monitoring the level of circulating EVs in the bloodstream might reflect tumour size. Indeed, GBM patients have increased vesiclemia^5^, while resection normalized these values^43^. Thus, the level of plasmatic EVs may assist in the clinical management of GBM patients, and/or to anticipate recurrence as a companion diagnostic tool. Overall, our work unveils the potential of interfering with EV biogenesis as new combined therapy to sensitize glioblastoma cell to death.

## Methods

### The Cancer Genome Atlas (TCGA) Analysis

The Cancer Genome Atlas (TCGA) database was explored via the Gliovis platform (http://gliovis.bioinfo.cnio.es)^32^. We interrogated RNAseq data from glioma patients for the expression of *MLKL* and EV-related genes, as well as probability of survival, according to histological glioma status (glioma grades II to IV), glioblastoma subtypes (classical, mesenchymal, neuronal and proneuronal) and *IDH1* mutation status.

### Cell culture and MLKL targeting

Patient-derived glioblastoma cells with stem-like properties (GSCs) were obtained from primary glioblastoma resection, as previously described^36^. Mesenchymal (MES1 and MES4) and classical (CL9) GSCs were maintained as spheres in NS34 medium (DMEM-F12 with N2, G5, and B27 supplements, plus glutamax and antibiotics, Life Technologies). Differentiated glioblastoma cells (DGCs) were generated from GSCs and expanded in DMEM/F12 with 10% foetal bovine serum (FBS), glutamax and antibiotics (Life Technologies) for at least 10 days. Differentiation was controlled using morphology, TUBB3 increase in expression, and SOX2 and NESTIN loss of expression.

To knockdown *MLKL* expression, RNA interference was performed using 10 pmol of duplexes targeting human *MLKL* (GCAACGCAUGCCUGUUUCACCCAUA, Stealth siRNA from Life Technologies) or non-silencing control duplexes (low-GC 12935111, Life Technologies) mixed with OptiMEM and Lipofectamine RNAi-MAX Transfection Reagent (Life Technologies) as previously described^31^. Alternatively, endogenous MLKL protein was depleted taking advantage of the Trim-away TRIM21/proteasome system recently described^40,65^. Briefly, MLKL antibody (Abcam ab211045) was electroporated in 5.10^5^ GSCs with Neon electroporation system according to manufacturer’s instructions (Life Technologies). Pharmacological targeting of MLKL was achieved using *in vitro* treatment with the MLKL inhibitor necrosulfonamide NSA (Abcam, 5 µM).

### Glioblastoma xenografts

Animal procedures were conducted in agreement with the European Convention for the Protection of Vertebrate Animals used for Experimental and other Scientific Purposes (ETS 123) and approved by the French Government (APAFIS#2016-2015092917067009). At all times, animals were allowed access to food and water *ad libitum* and were housed in specific pathogen free (SPF) environment with temperature and hygrometry controls on a 12-hour day-night cycle.

Ectopic and orthotopic glioblastoma models were adapted from previously described models^36^. CL9 GSCs were transduced with GFP and luciferase (pLNT-LucF/pFG12-eGFP mixture plasmid, a kind gift of Valérie Trichet, Université de Nantes, France). 10^6^ and 10^5^ modified patient-derived cells were implanted in Balb/c Nude (BALB/cAnNRj-Foxn1 nu/nu) mice (Janvier Labs), subcutaneously in each flank and orthotopically in the striatum, respectively, as previously described^36^. Tumour growth was monitored each week by bioluminescence on a PhotonIMAGER (Biospace Lab) 10min after injection of D-luciferin (Interchim FP-M1224D).

For toxicity experiments, NSA (5 mg/kg) was injected intraperitoneally in non-tumour bearing mice three times per week for 4 weeks. Toxicity score was determined each week depending on weight, body condition and mouse behaviour (grading from 1 for normal to 3 for critical). Liver transaminases ASAT and ALAT were measured at the hospital biology platform (CHU Hotel-Dieu, Nantes, France).

Five days post-GSC grafts, treatments with vehicle (PBS-DMSO) or NSA (5 mg/kg) was commenced three times a week, until critical point was reached or the conclusion of the experiment at day 50. At euthanasia, indicated organs were weighed and frozen; total blood was collected by intracardiac puncture on EDTA tubes and centrifugated (1000g, 15min, 4°C) before freezing at −80°C.

### Immunoblotting

For human tissues, protein lysates from normal brain and brain tumour were obtained from NovusBio and GeneTex companies. For murine tissues, brain biopsies were harvested at euthanasia and lysed in RIPA (25 mM Tris pH7.4, 150 mM NaCl, 0.1% SDS, 0.5% Na-Deoxycholate, 1% NP40, 1 mM EDTA, supplemented with protease inhibitor cocktail (Life Technologies) for 2h under agitation at 4°C following dispersion with pestle and mortar. Lysates were clarified by centrifugation (14000g, 30min) to remove debris. For cell culture, lysis was performed, after a wash with cold PBS, in TNT buffer (50 mM TRIS pH7.4, 150 mM NaCl, 1% Triton X100, 1% Igepal, 2 mM EDTA, supplemented with protease inhibitor cocktail) for 30 minutes on ice. Lysates were clarified at 10000g for 5min. Proteins concentration was determined using MicroBCA protein assay kit (Thermo Scientific). 10 µg of proteins were resolved in Tris-acetate SDS-PAGE and transferred to nitrocellulose membranes (GE Healthcare). For extracellular vesicle fractions, pelleted EVs from identical number of cells were directly lysed in boiling Laemmli 2X. Membranes were revealed using a chemiluminescent HRP substrate (Millipore) and visualised using the Fusion imaging system (Vilber Lourmat).

### Antibodies

The following primary antibodies were used: MLKL (Cell Signaling 14993), GAPDH (Santa Cruz sc-32233), ALIX (BioLegend 6345.02), CD9 (SBI EXOAB-CD9A-1), CD63 for WB (SBI EXOAB-CD63A-1), CD63 for immunofluorescence staining (BD 556019), eGFP (Millipore AB10145), SOX2 (Millipore AB5603), NESTIN (Millipore MAB5326), EEA1 (BD Bioscience 610456), GM130 (Abcam ab52649), Lamp2 (Santa Cruz sc-18822), RAB27A (Cell Signaling 95394), phospho-Histone H3 (Cell Signaling 33775), cleaved-caspase 3 (Cell Signaling 96645), phospho-ATM (Cell Signaling 5883). HRP-conjugated secondary antibodies (1/5000) were from Southern Biotech. Alexa-conjugated secondary antibodies were from Life Technologies.

### RT-PCR and qPCR

RNA extraction was done using Qiagen RNeasy kit and equal amounts of RNA were reverse transcribed using the Maxima Reverse Transcriptase kit (ThermoFisher). Semi-quantitative qPCR was performed using PerfeCTa SYBR Green SuperMix (QuantaBio). The following primers were used: human *MLKL* forward GCCACTGGAGATATCCCGTT, human *MLKL* reverse CTTCTCCCAGCTTCTTGTCC, human *SOX2* forward CAAAAATGGCCATGCAGGTT, human *SOX2* reverse AGTTGGGATCGAACAAAAGCTATT, human *TUBB3* forward CAGATGTTCGATGCCAAGAA, human *TUBB3* reverse GGGATCCACTCCACGAAGTA, human *ACTB* forward GGACTTCGAGCAAGAGATGG, human *ACTB* reverse AGCACTGTGTTGGCGTACAG, human *HPRT1* forward TGACACTGGCAAAACAATGCA, human *HPRT1* reverse GGTCCTTTTCACCAGCAAGCT. Data was analysed by the 2-ΔΔCt methods and normalised using the two housekeeping genes ACTB and HPRT1.

### Isolation of Extracellular Vesicles (EVs)

GSC-derived EVs were isolated from 4 to 5.10^6^ GSCs seeded in 10mL NS34 serum-free media for 48h. DGC-derived EVs were isolated from 80% confluent cells in 10mL DMEM/F12 with 10% foetal bovine serum (FBS), glutamax and antibiotics (Life Technologies) for 48h. Complete media was depleted from EVs with 16h-ultracentrifugation at 100,000g 4°C prior incubation with DGCs. Differential centrifugation was performed to isolate EVs from conditioned media: 300g 3min, 2000g 10min, 10000g 30min (10K fraction) and 100000g 2h (100K) fraction on a Beckman Coulter ultracentrifuge (OPTIMA L-80) using SW-41 Ti rotor. 10K and 100K pellets were washed in 0.22µm filtered PBS, ultracentrifuged (100000g, 2h) and resuspended in filtered PBS. OptiPrep top to bottom density gradients 5 % to 40 % were performed as described^66^ in 12mL open top polyallomer tubes (Beckman Coulter). 1mL fractions were collected from the top. To isolate murine circulating EVs, plasma separation from total blood harvested by intracardiac punction was performed at 1000g, 15min, and frozen at −80°C until further use. 200 µL of thawed plasmas were centrifuged at 2000g 10min, 10000g 30min and 100000g 2h. Experimental parameters were submitted to the open-source EV-TRACK knowledgebase (EV-TRACK.org)^67^, EV-TRACK ID is EV210024.

### Quantification of EVs

Single particle tracking was performed by Tunable Resistive Pulse Sensing (TRPS) using qNano gold apparatus (IZON) as preciously described^5^. Diameter and concentration of 100K and 10K EVs were determined using np100 nanopore (detected range 50−330 nm) and np400 (185-1100nm), respectively. Alternatively, abundance of EVs was determined using ELISA kit for CD63-positive particles according to supplier instructions (SBI). EV concentration was also determined using the newly developed interferometry light microscopy (Videodrop, Myriade) measuring size and concentration in a real-time nanometer scale optical method (interferometry).

### Electron microscopy

Electron microscopy images of EVs were acquired thanks to Joëlle Veziers (SC3M core-facility, CHU Nantes, France). Isolated EVs were fixed in glutaraldehyde 1.6%, diluted in 0.1M phosphate buffer, pH=7.2, coated on formvar carbon copper grids (Agar Scientific), stained with Uranyless (Delta Microscopies) and observed with a Jeol JEM-1010 transmission electron microscope. Relative EV size quantification (arbitrary units) was performed using Fiji surface area (n=380 and n=121 for sic *versus* si*MLKL*). Electron microscopy images were performed on GSCs. GSCs were fixed in 2.5% glutaraldehyde/2.0% paraformaldehyde, diluted in 0.1M Cacodylate buffer, at 4°C overnight. Samples were then stained in 0.05% Green malachite/2.5% glutaraldehyde and post-fixed with 1%OsO_4_ and 0.8% K_4_Fe(CN)_6_ for 45 min on ice. Samples were stained with 1% Tannic acid for 20 min on ice and with 1% uranyl acetate, overnight at 4°C. Samples were stepwise dehydrated in Ethanol and embedded in Epon. 100 nm thin sections were collected in cupper mesh grids, contrasted with 1% uranyl acetate followed with 0.4% lead citrate (Sigma-Aldrich) and imaged with a Hitachi 7500 transmission electron microscope equipped with Hamamastu camera C4742-51-12NR.

### Confocal and super resolution microscopy

Immunostaining of GSCs was performed as previously described^60^. Briefly, cells were adhered onto poly-lysine slides, fixed with 4% PFA and permeabilised with Triton X100. Primary antibodies were incubated overnight in PBS-BSA 4% at 4°C and AlexaFluor-conjugated secondary antibodies for 45min at room temperature. Samples were mounted in Prolong gold anti-fade mountant with DAPI (Life Technologies). Confocal images were acquired on confocal Nikon A1Rsi (Nikon Excellence Center, MicroPICell facility, Nantes). Super-resolution microscopy using structure illumination microscopy (SIM) images were acquired with a Nikon N-SIM (Nikon Excellence Center, MicroPICell facility, Nantes) using a 100x oil-immersion lens with a 1.49 aperture and reconstructed in 3D using the NIS-Element Software. All images were analysed using the Fiji software.

### Proteomic analysis of EVs

Mass Spectrometry was done in collaboration with 3P5 Proteom’IC facility (Université de Paris, Institut Cochin, Paris, France). Peptide contents from 100K EV fractions were lysed and denatured in SDS 4%, 50mM Tris pH=8 (5□min at 95□°C). Disulfide bridges were reduced (TCEP 20□mM) and subsequent free thiol groups were protected using chloroacetamide 50□mM. Proteins were digested overnight with trypsin and prepared with S-trap (https://www.protifi.com/s-trap/). Nano Liquid Chromatography with Tandem Mass Spectrometry analysis (nLC-MS/MS) was then performed, peptides were concentrated, separated and analysed with an Ultimate 3000 Rapid Separation liquid chromatographic system coupled to a Q-exactive mass spectrometer (Thermo Fisher Scientific). Raw data were processed with MaxQuant software to perform comparison of experimental MS/MS peptides fragmentation data with the Homo sapiens taxon of the Swiss-Prot Uniprot database. Identified proteins were processed through the STRING online database (https://string-db.org).

### Transcriptomic analysis

Cells were lysed and RNA content was analysed by 3’ Sequencing RNA Profiling at the Genomics and Bioinformatics Core Facility (GenoBiRD, Biogenouest, IFB, Nantes). Briefly, total RNA was isolated using the Qiagen RNeasy Mini Kit (ThermoFisher) and clean samples (DO 260/280 and 260/230>1.8) were submitted for quality control on a Bioanalyzer TapeStation (Agilent). RNA libraries were prepared and HiSeq sequencing was carried out (HiSeq 2500, Illumina). Data were demultiplexed and analysed with Illumina Bcl2fastQ2 software. Reads were aligned against human reference transcriptome (hg19) and differential analysis was performed with DESeq2 tool and annotated with Gene Ontology and Kegg pathways (methods available on https://bio.tools/3SRP).

### Stemness assays

To analyse self-renewal properties of GSCs, tumoursphere formation and limiting dilution assay (LDA) were performed. For tumoursphere formation, GSCs (100/µL) were seeded in triplicate in NS34 media as previously described^36,60^. Cells were dissociated manually both the second and third days to reduce cell aggregation influence. At day 5, the number of tumourspheres per field of view (NS/fov) was calculated by counting 5 random fov per condition.

To analyse further self-renewal of GSCs, LDA was performed as previously described^36,60^, GSCs were plated in a 96-well plate using serial dilution ranging from 2000 to 1 cell per well (one column, *i*.*e*. 8 replicates per dilution) and treated or not with NSA. After 14 days of cell culture, each well was binary evaluated for the presence or the absence of tumoursphere. Stemness frequency was then calculated using online ELDA software (http://bioinf.wehi.edu.au/software/elda)^68^. The mean stemness frequency was calculated by averaging 3 independent experiments.

### Proliferation and cytotoxicity assays

GSC viability was assessed using Uptiblue (Interchim) and CellTiter-Glo (Promega) following manufacturers’ protocol. Absorbance and luminescence were measured on a FluoStarOptima (BMG Labtech) plate reader and the percentage of cell viability in comparison to control/vehicle conditions was determined.

For EdU proliferation analysis, cells were incubated with EdU (10 µM) for 2h followed by fixation and Click-it reaction according to instructions for flow cytometry (Life Technologies, C10424) or by secondary antibody incubation for confocal analysis (Life Technologies, C10337). Propidium Iodide (PI) and Annexin V staining (Life Technologies) were used to assess cell death induction following treatment according to manufacturer’s instructions.

### Flow cytometry

Flow Cytometry analyses were performed on FACSCalibur (BD Biosciences, CytoCell facility, Nantes) and processed using FlowJo software (BD Biosciences).

### Statistics

Data are expressed as mean ±SEM and are representative of at least three independent experiments, unless otherwise stated. Statistical analysis was performed with GraphPad Prism8 using two-way ANOVA, parametric *t*-test or non-parametric Mann-Whitney when required. For each statistical test, a p-value <0.05 was considered as significant.

## Acknowledgements

We are grateful to past and present SOAP team members, especially Sarah Delalleau (Université de Nantes, INSERM, CNRS, France). We also thank Francois Guillonneau, Cédric Broussard and Virginie Salnot from Plateforme Proteom’IC 3P5, Universite de Paris, Institut Cochin, Paris, France for performing sample preparation and data acquisition and analysis, Joëlle Veziers (SC3M, CHU Nantes, France) for image acquisition, Valérie Trichet (PhyOs UMR1238, Nantes, France) for eGFP-Luc plasmid and the Genomics and Bioinformatics Core Facility of Nantes (GenoBiRD, Biogenouest, IFB) for their technical assistance with 3’SRP analysis. We would like to acknowledge the core-facilities from SFR Santé François Bonamy, Nantes, France (MicroPICell, Cytocell, Therassay, and UTE IRS-UN) and Cathy Royer from the plateforme d’imagerie In Vitro, CNRS UPS3156, Strasbourg, France.

This work was supported by Fondation de France (GAG), Fondation pour la Recherche Médicale (Equipe labellisée DEQ20180339184), Fondation ARC contre le Cancer (JG, NB), INCa PLBIO (2019-151, 2019-291), Ligue nationale contre le cancer comités de Loire-Atlantique, Maine et Loire, Vendée, Ille-et-Vilaine (JG, NB). TD received a fellowship from Nantes Métropole, AT from Fondation ARC, KAJ from Nantes Métropole and Fondation ARC, CM from Ligue Régionale contre le Cancer. The team is part of the SIRIC ILIAD (INCA-DGOS-Inserm_12558).

## Author contributions

GAG, JG, NB, conception and design, acquisition of data, analysis and interpretation of data, drafting the article. TD, AT, KJ, CM, KT, CB, IB, VH, JGG acquisition of data, analysis and interpretation of data. All authors approved the manuscript.

## Supplemental Figure Legends

**Supplementary Figure 1.**
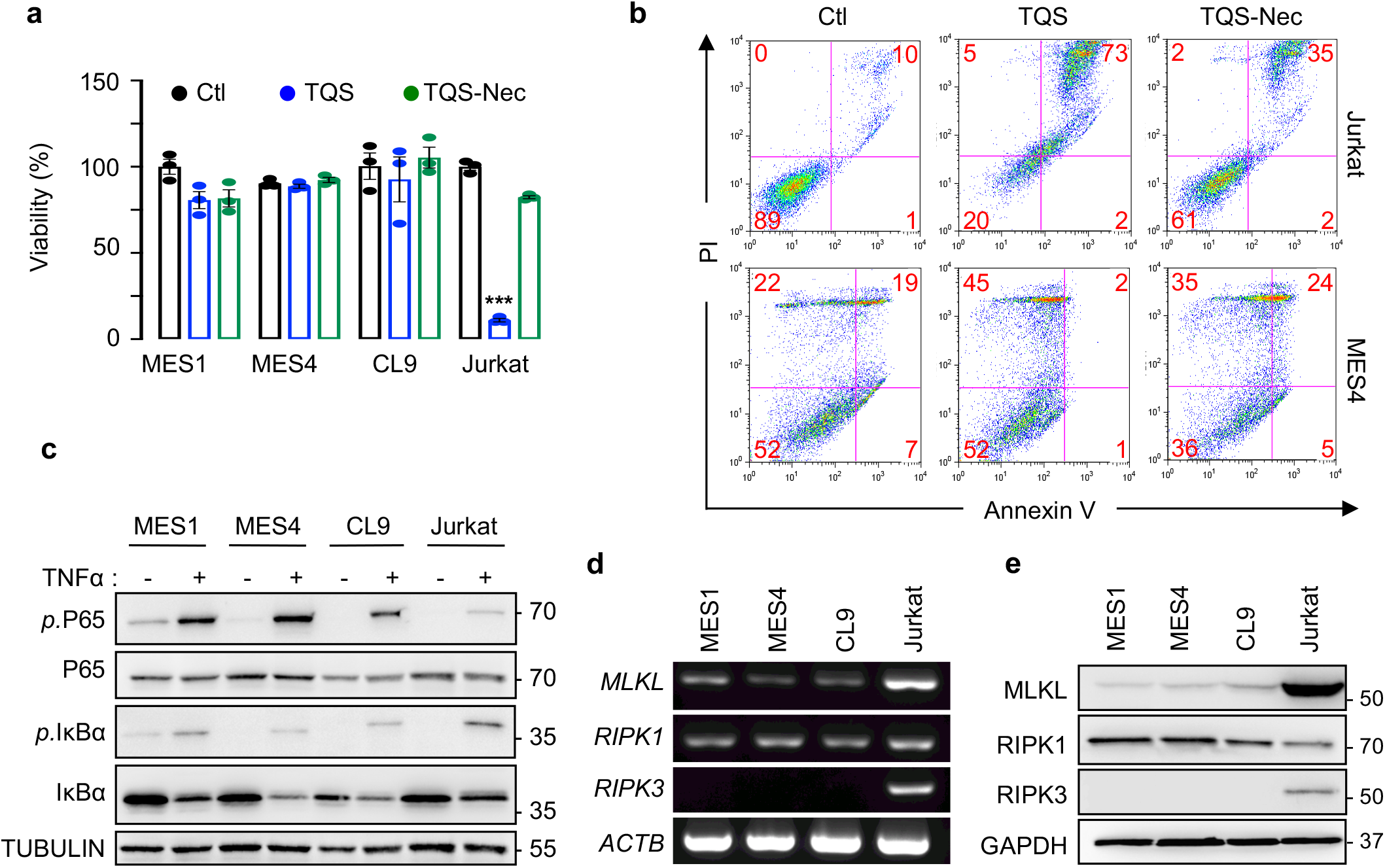
Glioblastoma Stem-like Cells are resistant to necroptosis. **a** Uptiblue viability assay in MES1, MES4, CL9 and Jurkat upon necroptosis stimulation with TNFα (T, 10ng.mL-1), caspase inhibitor QVD (Q, 10 µM), and the SMAC-mimetic Birinapant (S, 5 µM), in the presence or absence of the necroptosis inhibitor Necrostatin-1 (Nec, 20 µM), n=3. **b** Representative flow cytometry analysis of propidium iodide (PI) and Annexin V staining in MES4 and Jurkat upon necroptosis stimulation with TQS, in the presence or absence of Necrostatin-1 (Nec, 20 µM), n=3. **c** Mesenchymal (MES) and classical (CL) Glioblastoma Stem-like Cells MES1, MES4 and CL9 were stimulated with TNFα (10ng.mL-1, 15 min). Protein lysates were analysed by immunoblotting, with the indicated antibodies (*p*=phosphorylation). Jurkat T lymphocytes serve as a positive control. TUBULIN serves a loading control. **d** mRNA of *RIPK1, RIPK3* and *MLKL* were detected in indicated cells by RT-PCR and representative agarose gel electrophoresis is shown, n=2. *ACTB* serves as internal control. **e** Immunoblot analysis of RIPK1, RIPK3 and MLKL protein expression in MES1, MES4, CL9, and Jurkat, n=3. GAPDH serves a loading control. Data are representative of at least three independent experiments, unlike otherwise stated. One-way ANOVA: *** p <0.001.

**Supplementary Figure 2.**
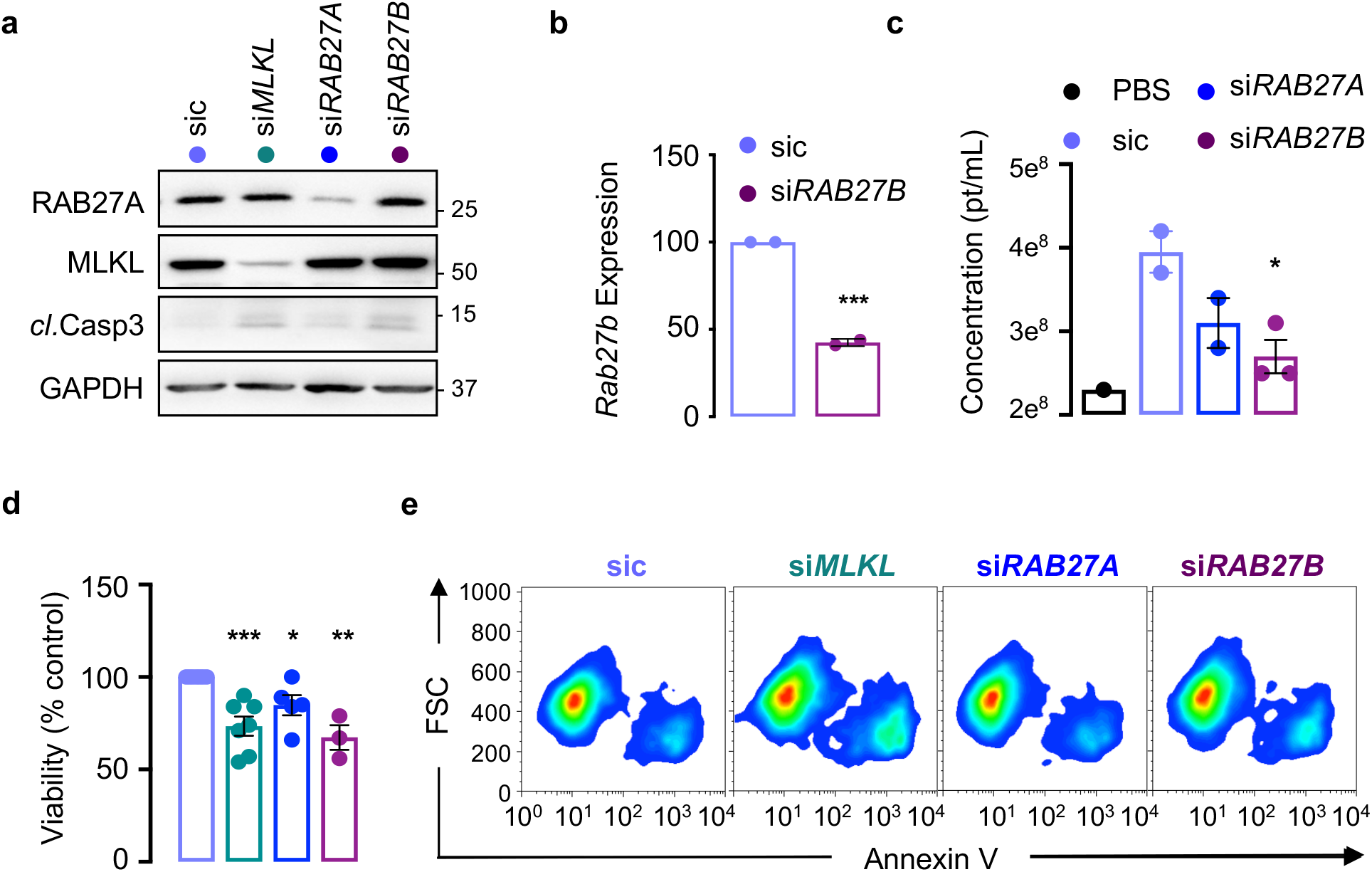
Rab27a-b deficiency mimics MLKL impairment in GSCs. **a** Mesenchymal (MES) Glioblastoma Stem-like Cells MES4 were transfected with non-silencing (sic), *MLKL, RAB27A* and *RAB27B* RNA duplexes, as indicated. Protein lysates were harvested 72h later and analysed by immunoblot for MLKL, RAB27A, and Cleaved caspase-3 (*cl*.Casp3), n>2. GAPDH serves a loading control. **b** mRNA Expression of *RAB27B* was detected by RT-qPCR analysis in sic and si*RAB27B* transfected MES4, n=2. **c** 100k small EVs were isolated from the 48h-old conditioned media of serum-free, EV-free of transfected sic, si*MLKL*, si*RAB27A* and si*RAB27B* transfected MES4, and quantified by interferometry light microscopy (Videodrop, Myriade), n>2. **d** Uptiblue viability assay from MES4 as in panel (c), n>3. **e** Representative flow cytometry analysis of the cell death marker Annexin V surface staining in MES4 treated as in panel (a), n>2. FSC: Forward scatter. Data are representative of at least three independent experiments, unlike otherwise stated. t-test and one-way ANOVA, *p<0.05, **p<0.01, ***p<0.001.

**Supplementary Figure 3.**
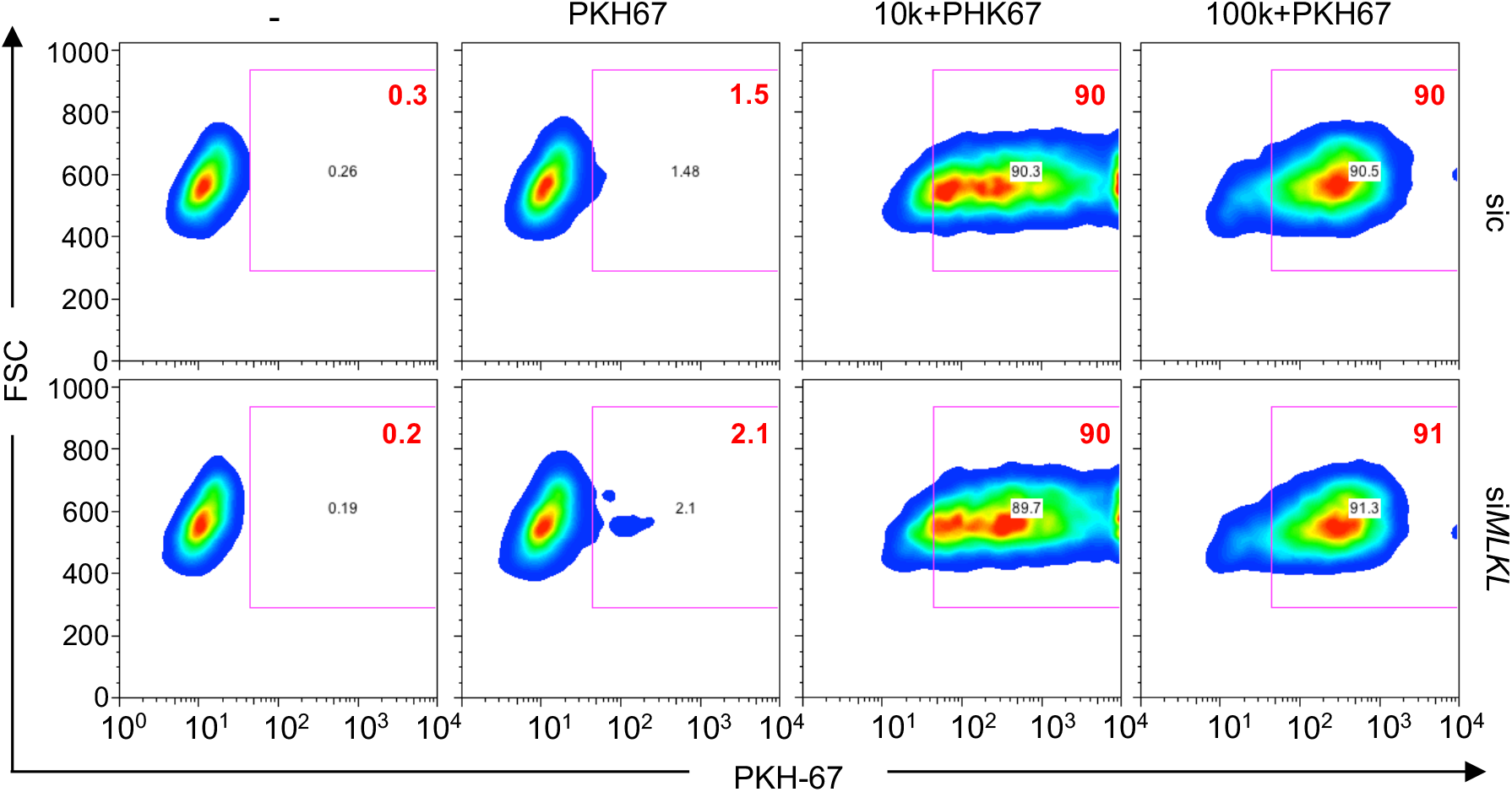
Extracellular Vesicles uptake and rescue. Small (100k) and large (10k) EVs were isolated from mesenchymal Glioblastoma Stem-like Cells (MES4 GSCs) and further stained with the lipidic probe PKH67. Representative flow cytometry analysis of PKH67-stained EV uptake in classical GSCs (CL9) after 4h, n>2. As a control, CL9 received PBS (-) and PKH67-stained cell-free 100k fraction (PKH67). Data are representative of at least three independent experiments.

**Supplementary Figure 4.**
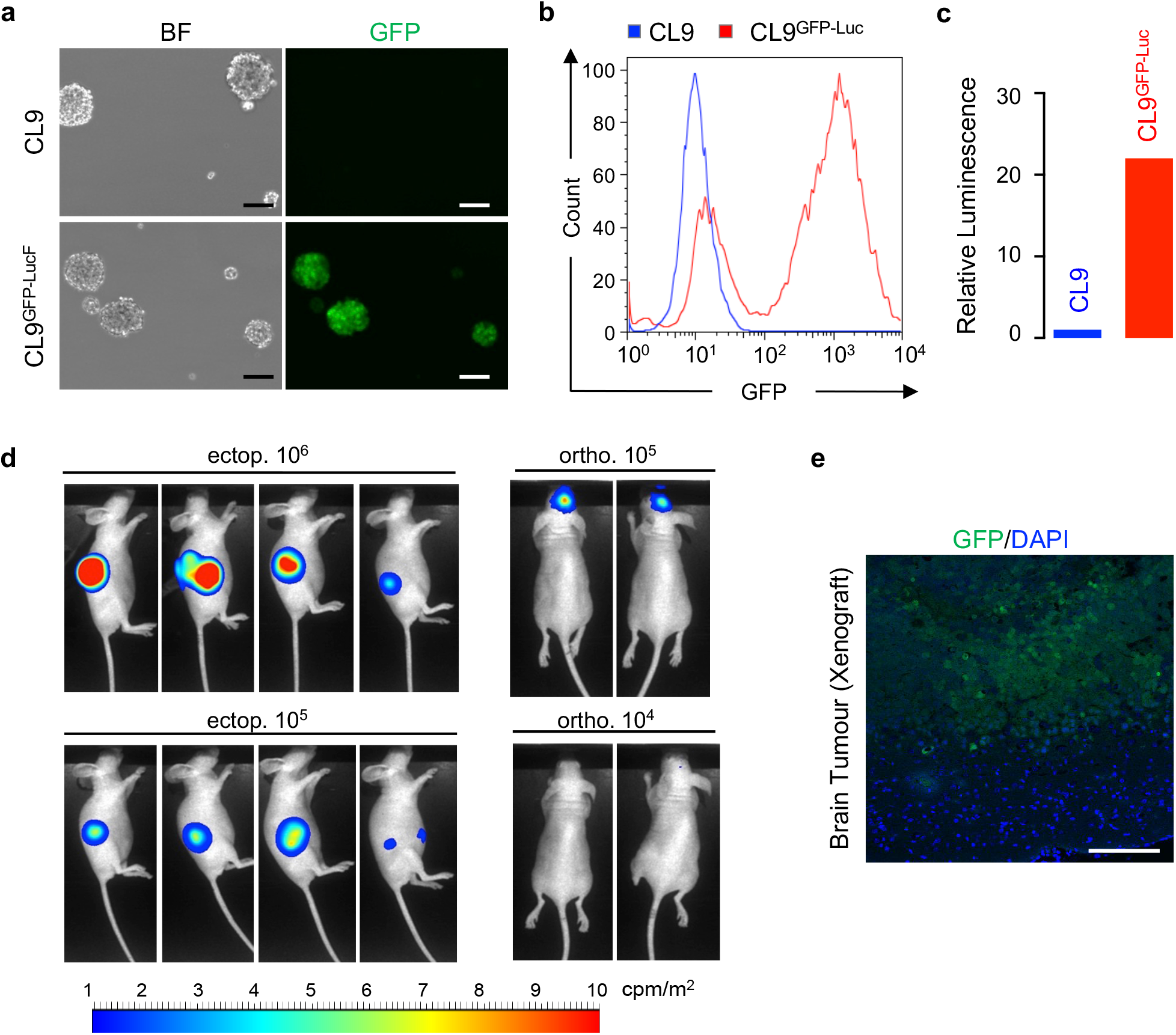
Preparation of modified patient-derived GSC for GBM xenografts in mice. **a** Representative bright field (BF) and fluorescent images (GFP) of patient-derived classical Glioblastoma Stem-like Cells (CL9 GSCs) before and after transduction with eGFP/luc. Scale bar=100 μm. **b** Representative flow cytometry analysis of GFP fluorescence in CL9 and CL9^GFP-Luc^ cells. **c** Relative luminescence from CL9 and CL9^GFP-Luc^ cell lysates. **d** 10^6^ and 10^5^ CL9^GFP-Luc^ cells were implanted in nude mice ectopically (subcutaneous, ectop.) and orthotopically (brain, ortho.), respectively. **e** Representative confocal image of xenografts in mouse brains. Nuclei are counterstained with DAPI (blue). Scale bar=100 μm.

**Supplementary Table 1. Proteomic analysis of EVs isolated from NSA-treated GSCs.**

List of proteins identified by mass spectrometry analysis (LC-MS/MS) of small (100k) EVs isolated from mesenchymal Glioblastoma Stem-like Cells (MES4 GSCs) treated with vehicle (DMSO) or MLKL inhibitor necrosulfonamide (NSA, 5 μM, 48 hours).

**Supplementary Table 2. 3’SRP transcriptomic analysis of NSA-treated GSCs.**

List of differentially expressed genes (deseq2) in mesenchymal Glioblastoma Stem-like Cells MES4 treated with vehicle (DMSO) or necrosulfonamide (NSA, 5 μM, 16 or 48 hours) and analysed through 3’Seq RNA Profiling.

**Supplementary Table 3. Blood count in mice challenged with NSA.**

For evaluation of NSA tolerance/toxicity *in vivo* in mice, blood count analysis (Hemavet) in EDTA-plasma from vehicle and NSA treated mice for 4 weeks, n=2 (vehicle) and 3 (NSA).

**Supplementary Movie 1.**

Real-time recording of brightfield classical Glioblastoma Stem-like Cells CL9 during 4 days of DMSO or NSA treatment (5 μM) using the IncuCyte live-cell imaging system.

## Supplemental information

### Cell culture and chemicals

T-cell Jurkat were purchased from American Type Culture Collection. Necroptosis was induced *in vitro* by pre-treatment with Q-VD-OPh (R&D System, 10 µM) and Birinapant (Selleckchem, 5 µM) prior stimulation with TNFα (R&D Systems, 10ng.mL-1). Necroptosis was blocked using Necrostatin Nec-1s (Selleckchem, 20 µM).

### Antibodies

The following primary antibodies were used: phospho-p65 (Cell Signaling 3033), P65 (Santa Cruz sc-372), phospho-IκBα (Cell Signaling 9246), IκBα (Cell Signaling 4814), RIPK1 (BD 551042), RIPK3 (Bethyl 13526), TUBULIN (Santa Cruz sc-8035).

### RT-PCR and qPCR

Classical PCR was performed on the resulting cDNA using Red Taq DNA polymerase (Sigma) or semi-quantitative qPCR, as described in the manuscript. RIPK1 forward AGTGACTTCCTGGAGAGTGC, RIPK1 reverse TCATCATCTTCGCCTCCTCC, RIPK3 forward CTCTCTGCGAAAGGACCAAG, RIPK3 reverse TCGTAGCCCCACTTCCTATG, RAB27B forward CTTCGCAGGCTGACCGA, RAB27B reverse CCACACACTGTTCCATTCGC.

### siRNA

RNA interference with *RAB27A* and *RAB27B* was performed as previously described^31^.

### Uptake of EVs

PKH-67 staining of EVs was performed as previously described^13^. 5×10^5^ GSCs were seeded in 1 mL of NS34 with the labelled EVs for 4 hours. After 10min fixation in 4% PFA and washes, flow cytometry analyses were performed on a FACSCalibur (BD Biosciences, CytoCell facility, Nantes) and processed using FlowJo software.

### Incucyte

Real-time effect of NSA treatment was recorded on alive cells using IncuCyte Zoom live cell imaging system (Essen BioScience). Briefly, GSCs were seeded at a density of 5000/well in a 96-well plate, treated with NSA (Abcam, 5 µM) or vehicle and maintained at 37°C and 5% CO2 during the time of experiment. Phase contrast images (10X zoom) were taken every 2 hours for 4 days.

## References

1. Stupp, R. et al.. Radiotherapy plus concomitant and adjuvant temozolomide for glioblastoma. N. Engl. J. Med. 352, 987–996 (2005).

2. Gimple, R. C., Bhargava, S., Dixit, D. & Rich, J. N. Glioblastoma stem cells: lessons from the tumor hierarchy in a lethal cancer. Genes Dev. 33, 591–609 (2019).

3. Bao, S. et al.. Glioma stem cells promote radioresistance by preferential activation of the DNA damage response. Nature 444, 756–760 (2006).

4. Chen, J. et al.. A restricted cell population propagates glioblastoma growth after chemotherapy. Nature 488, 522–526 (2012).

5. André-Grégoire, G., Bidère, N. & Gavard, J. Temozolomide affects Extracellular Vesicles Released by Glioblastoma Cells. Biochimie 155, 11–15 (2018).

6. André-Grégoire, G. & Gavard, J. Spitting out the demons: Extracellular vesicles in glioblastoma. Cell Adhes. Migr. 11, 164–172 (2017).

7. Quezada, C. et al.. Role of extracellular vesicles in glioma progression. Mol. Aspects Med. 60, 38–51 (2018).

8. van Niel, G., D’Angelo, G. & Raposo, G. Shedding light on the cell biology of extracellular vesicles. Nat. Rev. Mol. Cell Biol. 19, 213–228 (2018).

9. Gao, X. et al.. Gliomas Interact with Non-glioma Brain Cells via Extracellular Vesicles. Cell Rep. 30, 2489-2500.e5 (2020).

10. Al-Nedawi, K. et al.. Intercellular transfer of the oncogenic receptor EGFRvIII by microvesicles derived from tumour cells. Nat. Cell Biol. 10, 619–624 (2008).

11. Luhtala, N., Aslanian, A., Yates, J. R. & Hunter, T. Secreted Glioblastoma Nanovesicles Contain Intracellular Signaling Proteins and Active Ras Incorporated in a Farnesylation-dependent Manner. J. Biol. Chem. 292, 611–628 (2017).

12. Skog, J. et al.. Glioblastoma microvesicles transport RNA and proteins that promote tumour growth and provide diagnostic biomarkers. Nat. Cell Biol. 10, 1470–1476 (2008).

13. Treps, L. et al.. Extracellular vesicle-transported Semaphorin3A promotes vascular permeability in glioblastoma. Oncogene 35, 2615–2623 (2016).

14. Treps, L., Perret, R., Edmond, S., Ricard, D. & Gavard, J. Glioblastoma stem-like cells secrete the pro-angiogenic VEGF-A factor in extracellular vesicles. J. Extracell. Vesicles 6, 1359479 (2017).

15. de Vrij, J. et al.. Glioblastoma-derived extracellular vesicles modify the phenotype of monocytic cells. Int. J. Cancer 137, 1630–1642 (2015).

16. Harshyne, L. A., Nasca, B. J., Kenyon, L. C., Andrews, D. W. & Hooper, D. C. Serum exosomes and cytokines promote a T-helper cell type 2 environment in the peripheral blood of glioblastoma patients. Neuro-Oncol. 18, 206–215 (2016).

17. Garnier, D. et al.. Divergent evolution of temozolomide resistance in glioblastoma stem cells is reflected in extracellular vesicles and coupled with radiosensitization. Neuro-Oncol. 20, 236–248 (2018).

18. Yin, J. et al.. Exosomal transfer of miR-1238 contributes to temozolomide-resistance in glioblastoma. EBioMedicine 42, 238–251 (2019).

19. Zeng, A. et al.. Exosomal transfer of miR-151a enhances chemosensitivity to temozolomide in drug-resistant glioblastoma. Cancer Lett. 436, 10–21 (2018).

20. Zhang, Z. et al.. Exosomal transfer of long non-coding RNA SBF2-AS1 enhances chemoresistance to temozolomide in glioblastoma. J. Exp. Clin. Cancer Res. CR 38, 166 (2019).

21. Zhao, J. et al.. Mixed lineage kinase domain-like is a key receptor interacting protein 3 downstream component of TNF-induced necrosis. Proc. Natl. Acad. Sci. U. S. A. 109, 5322–5327 (2012).

22. Sun, L. et al.. Mixed lineage kinase domain-like protein mediates necrosis signaling downstream of RIP3 kinase. Cell 148, 213–227 (2012).

23. Dai, J. et al.. A necroptotic-independent function of MLKL in regulating endothelial cell adhesion molecule expression. Cell Death Dis. 11, 282 (2020).

24. Ying, Z. et al.. Mixed Lineage Kinase Domain-like Protein MLKL Breaks Down Myelin following Nerve Injury. Mol. Cell 72, 457-468.e5 (2018).

25. Kang, S. et al.. Caspase-8 scaffolding function and MLKL regulate NLRP3 inflammasome activation downstream of TLR3. Nat. Commun. 6, 7515 (2015).

26. Conos, S. A. et al.. Active MLKL triggers the NLRP3 inflammasome in a cell-intrinsic manner. Proc. Natl. Acad. Sci. U. S. A. 114, E961–E969 (2017).

27. Gong, Y.-N. et al.. ESCRT-III Acts Downstream of MLKL to Regulate Necroptotic Cell Death and Its Consequences. Cell 169, 286-300.e16 (2017).

28. Yoon, S., Kovalenko, A., Bogdanov, K. & Wallach, D. MLKL, the Protein that Mediates Necroptosis, Also Regulates Endosomal Trafficking and Extracellular Vesicle Generation. Immunity 47, 51-65.e7 (2017).

29. Zargarian, S. et al.. Phosphatidylserine externalization, ‘necroptotic bodies’ release, and phagocytosis during necroptosis. PLoS Biol. 15, e2002711 (2017).

30. Shlomovitz, I. et al.. Proteomic analysis of necroptotic extracellular vesicles. bioRxiv 2020.04.11.037192 (2020) doi:10.1101/2020.04.11.037192.

31. Douanne, T. et al.. Pannexin-1 limits the production of proinflammatory cytokines during necroptosis. EMBO Rep. 20, e47840 (2019).

32. Bowman, R. L., Wang, Q., Carro, A., Verhaak, R. G. W. & Squatrito, M. GlioVis data portal for visualization and analysis of brain tumor expression datasets. Neuro-Oncol. 19, 139–141 (2017).

33. Yan, H. et al.. IDH1 and IDH2 mutations in gliomas. N. Engl. J. Med. 360, 765–773 (2009).

34. Verhaak, R. G. W. et al.. Integrated genomic analysis identifies clinically relevant subtypes of glioblastoma characterized by abnormalities in PDGFRA, IDH1, EGFR, and NF1. Cancer Cell 17, 98–110 (2010).

35. Singh, S. K. et al.. Identification of human brain tumour initiating cells. Nature 432, 396–401 (2004).

36. Harford-Wright, E. et al.. Pharmacological targeting of apelin impairs glioblastoma growth. Brain J. Neurol. 140, 2939–2954 (2017).

37. Théry, C., Amigorena, S., Raposo, G. & Clayton, A. Isolation and Characterization of Exosomes from Cell Culture Supernatants and Biological Fluids. Curr. Protoc. Cell Biol. 30, 3.22.1-3.22.29 (2006).

38. Théry, C. et al.. Minimal information for studies of extracellular vesicles 2018 (MISEV2018): a position statement of the International Society for Extracellular Vesicles and update of the MISEV2014 guidelines. J. Extracell. Vesicles 7, 1535750 (2018).

39. Ostrowski, M. et al.. Rab27a and Rab27b control different steps of the exosome secretion pathway. Nat. Cell Biol. 12, 19–30; sup pp 1-13 (2010).

40. Clift, D. et al.. A Method for the Acute and Rapid Degradation of Endogenous Proteins. Cell 171, 1692-1706.e18 (2017).

41. Caporali, S. et al.. DNA damage induced by temozolomide signals to both ATM and ATR: role of the mismatch repair system. Mol. Pharmacol. 66, 478–491 (2004).

42. Lenting, K., Verhaak, R., ter Laan, M., Wesseling, P. & Leenders, W. Glioma: experimental models and reality. Acta Neuropathol. (Berl.) 133, 263–282 (2017).

43. Osti, D. et al.. Clinical Significance of Extracellular Vesicles in Plasma from Glioblastoma Patients. Clin. Cancer Res. Off. J. Am. Assoc. Cancer Res. 25, 266–276 (2019).

44. Sabbagh, Q., Andre-Gregoire, G., Guevel, L. & Gavard, J. Vesiclemia: counting on extracellular vesicles for glioblastoma patients. Oncogene 39, 6043–6052 (2020).

45. Murphy, J. M. et al.. The pseudokinase MLKL mediates necroptosis via a molecular switch mechanism. Immunity 39, 443–453 (2013).

46. Cai, Z. et al.. Plasma membrane translocation of trimerized MLKL protein is required for TNF-induced necroptosis. Nat. Cell Biol. 16, 55–65 (2014).

47. Yu, J. et al.. 2-Methoxy-6-acetyl-7-methyljuglone (MAM) induced programmed necrosis in glioblastoma by targeting NAD(P)H: Quinone oxidoreductase 1 (NQO1). Free Radic. Biol. Med. 152, 336–347 (2020).

48. Zhou, J. et al.. Emodin induced necroptosis in the glioma cell line U251 via the TNF-α/RIP1/RIP3 pathway. Invest. New Drugs 38, 50–59 (2020).

49. Melo-Lima, S., Celeste Lopes, M. & Mollinedo, F. Necroptosis is associated with low procaspase-8 and active RIPK1 and −3 in human glioma cells. Oncoscience 1, 649–664 (2014).

50. Miki, Y., Akimoto, J., Moritake, K., Hironaka, C. & Fujiwara, Y. Photodynamic therapy using talaporfin sodium induces concentration-dependent programmed necroptosis in human glioblastoma T98G cells. Lasers Med. Sci. 30, 1739–1745 (2015).

51. Koo, G.-B. et al.. Methylation-dependent loss of RIP3 expression in cancer represses programmed necrosis in response to chemotherapeutics. Cell Res. 25, 707–725 (2015).

52. Wang, Q. et al.. Epigenetic Regulation of RIP3 Suppresses Necroptosis and Increases Resistance to Chemotherapy in NonSmall Cell Lung Cancer. Transl. Oncol. 13, 372–382 (2020).

53. Moriwaki, K., Bertin, J., Gough, P. J., Orlowski, G. M. & Chan, F. K. M. Differential roles of RIPK1 and RIPK3 in TNF-induced necroptosis and chemotherapeutic agent-induced cell death. Cell Death Dis. 6, e1636 (2015).

54. Sun, Y. et al.. Down-regulation of RIP3 potentiates cisplatin chemoresistance by triggering HSP90-ERK pathway mediated DNA repair in esophageal squamous cell carcinoma. Cancer Lett. 418, 97–108 (2018).

55. Vergara, G. A., Eugenio, G. C., Malheiros, S. M. F., Victor, E. da S. & Weinlich, R. RIPK3 is a novel prognostic marker for lower grade glioma and further enriches IDH mutational status subgrouping. J. Neurooncol. 147, 587–594 (2020).

56. Bobrie, A. et al.. Rab27a supports exosome-dependent and -independent mechanisms that modify the tumor microenvironment and can promote tumor progression. Cancer Res. 72, 4920–4930 (2012).

57. van Solinge, T. S. et al.. Versatile Role of Rab27a in Glioma: Effects on Release of Extracellular Vesicles, Cell Viability, and Tumor Progression. Front. Mol. Biosci. 7, 554649 (2020).

58. Liu, Y., Zhou, Y. & Zhu, K. Inhibition of glioma cell lysosome exocytosis inhibits glioma invasion. PloS One 7, e45910 (2012).

59. Shingu, T. et al.. Qki deficiency maintains stemness of glioma stem cells in suboptimal environment by downregulating endolysosomal degradation. Nat. Genet. 49, 75–86 (2017).

60. Jacobs, K. A. et al.. Paracaspase MALT1 regulates glioma cell survival by controlling endo-lysosome homeostasis. EMBO J. 39, e102030 (2020).

61. Wu, J. et al.. Mlkl knockout mice demonstrate the indispensable role of Mlkl in necroptosis. Cell Res. 23, 994–1006 (2013).

62. Rathkey, J. K. et al.. Chemical disruption of the pyroptotic pore-forming protein gasdermin D inhibits inflammatory cell death and sepsis. Sci. Immunol. 3, (2018).

63. Bartzatt, R., Cirillo, S. L. G. & Cirillo, J. D. Sulfonamide agents for treatment of Staphylococcus MRSA and MSSA infections of the central nervous system. Cent. Nerv. Syst. Agents Med. Chem. 10, 84–90 (2010).

64. Bronisz, A. et al.. Extracellular Vesicles Modulate the Glioblastoma Microenvironment via a Tumor Suppression Signaling Network Directed by miR-1. Cancer Res. 74, 738–750 (2014).

65. Clift, D., So, C., McEwan, W. A., James, L. C. & Schuh, M. Acute and rapid degradation of endogenous proteins by Trim-Away. Nat. Protoc. 13, 2149–2175 (2018).

66. Van Deun, J. et al.. The impact of disparate isolation methods for extracellular vesicles on downstream RNA profiling. J. Extracell. Vesicles 3, (2014).

67. EV-TRACK Consortium et al. EV-TRACK: transparent reporting and centralizing knowledge in extracellular vesicle research. Nat. Methods 14, 228–232 (2017).

68. Hu, Y. & Smyth, G. K. ELDA: extreme limiting dilution analysis for comparing depleted and enriched populations in stem cell and other assays. J. Immunol. Methods 347, 70–78 (2009).

